# Chromodomain protein regulates the expression of a subset of RIFINs in *P. falciparum*

**DOI:** 10.1101/2021.09.13.460021

**Authors:** Devadathan Valiyamangalath Sethumadhavan, Marta Tiburcio, Abhishek Kanyal, CA Jabeena, Gayathri Govindaraju, Krishanpal Karmodiya, Arumugam Rajavelu

## Abstract

*Plasmodium falciparum* expresses clonally variant proteins on the surface of infected erythrocytes to evade the host immune system. The clonally variant multigenes include *var, rifin*, and *stevor*, which express EMP1, RIFIN, and STEVOR proteins, respectively. The *rifins* are the largest multigene family and are essentially involved in the RBC rosetting, the hallmark of severe malaria. The regulators that control the RIFINs expression in *P. falciparum* have not been reported so far. This study reports a chromodomain-containing protein (PfCDP) that binds to H3K9me3 modification on *P. falciparum* chromatin. The ChIP-sequencing analysis revealed that the PfCDP is majorly associated with clonally variant gene families, primarily *rifins* in *P. falciparum*. Conditional deletion of the chromodomain (CD) gene in *P. falciparum* leads to the up-regulation of a subset of virulence genes, including *rifins*, a few *var,* and *stevor* genes. Further, we show that PfΔCDP *P. falciparum* lines promote the RBC rosetting. This study provides evidence of an epigenetic regulator mediated control on a subset of RIFINs expression and RBC rosetting by *P. falciparum*.

## Introduction

Malaria remains a disease of global health importance, leading to an estimated 229 million cases and around 0.4 million deaths worldwide annually (WHO, 2019). Among malaria-causing parasites, *Plasmodium falciparum* causes the most severe form of malaria and is responsible for most malaria-associated human deaths (1, 2). The major clinical manifestations of the disease are associated with the erythrocytic stages of the infection. During this intra-erythrocytic developmental cycle (IDC), the parasite progresses through three distinct phases of development: ring, trophozoite, and schizont and the parasite relies on complex, poorly understood mechanisms to regulate the gene expression across these phases (3, 4). The emerging reports have identified that the epigenetic modifications contribute significantly to the gene regulation and are thus indispensable for the growth and development of *P. falciparum* (2, 5–11). The *Plasmodium spp.* contains unique epigenetic features such as the absence of linker histone H1 in their genome; it also lacks a functional RNA interference mechanism (12, 13). Although the presence of residual 5mC, 5hmC like modifications and methylation of tRNA Asp by DNA methyltransferase homolog were reported in *P. falciparum*, their involvement in gene regulation is poorly understood (14–16). These reports suggest that epigenetic mechanisms, mainly histone modifications might act as key players in regulating the gene expression in *Plasmodium spp*.

During the IDC of *P. falciparum*, several parasite-derived antigens are expressed on the surface of infected erythrocytes. These surface antigens, often known as variant surface antigens (VSA), are primary targets of the host immune defence mechanisms that include P. falciparum Erythrocyte Membrane Protein 1 (PfEMP1), Repetitive Interspersed Families of polypeptides (RIFINa), Sub telomeric Variable Open Reading frame (STEVOR), and surface associated interspersed protein (SURFIN) (17–20). These VSAs at the infected RBC (iRBC) surface facilitate the parasite’s escape from the host immune clearance and establish chronic infections (21). The PfEMP1, RIFINs, and STEVOR polypeptides are adhesive, and mediate the attachment of infected RBCs to the uninfected RBC (rosetting), endothelial cells (cytoadherence), leukocytes and serum proteins (22). It is known that the expression of these allelic variant genes is regulated by epigenetic chromatin modifications in *P. falciparum* (23–26).

The *P. falciparum* chromatin is enriched with modifications that are dynamically deposited/removed on the histones during its development in RBCs (2). Of these, the H3K9me3 and H3K36me3 marks are majorly found in repressive clusters containing the virulence family genes, such as *var, rifin, stevor, and pfmc-2tm* (23, 25). Apart from these virulence gene clusters, H3K9me3 is also present at several other loci, including invasion gene families (*eba* and *clag*) and pfap2-g (gametocyte-specific transcription factor) during the erythrocytic stages. The functions of these epigenetic methylation marks on the histone proteins are interpreted by specialized methyl recognition domain-containing proteins (27). The first epigenetic methyl-reader protein to be characterized in *Plasmodium spp*. was chromodomain containing PfHP1, which interacts with the H3K9me3 mark and essentially mediates the formation of heterochromatin and plays a crucial role in the regulation of *var* gene expression in *P. falciparum* (23, 24, 26, 28). A recent study has identified many putative methyl-reader complexes from *P. falciparum* using peptide pull-down assay followed by quantitative proteomics (29). Moreover, the chromodomain protein that we identified and characterized in this study had been reported in a recent study (29) to be one of the reader proteins of the H3K9me3 mark. However, its interaction with the H3K9me3 mark and its functional relevance in *P. falciparum* has not been addressed.

RIFINs are the vital contributors to immune evasion as they assist the rosetting of RBCs and also inhibit the immune cell (particularly B cell and NK cells) activation and leads to the development of severe malaria (30, 31). It is reported that nearly 185 copies of the *rifin* genes are present in the parasite genome. However, it is still unclear how the *rifin* gene expression is regulated and how many *rifins* are expressed by *P. falciparum* during the infection (32). Earlier reports have shown the enrichment of H3K9me3 repressor methylation marks on all suppressed *rifin* genes (23, 28); however, it is still unknown how the H3K9me3 mediates the suppression of the *rifin* genes in *P. falciparum*. Here, we report a chromodomain-containing protein (PfCDP) (PF3D7_1140700) that interacts with the H3K9me3 mark associated with virulence genes of *P. falciparum*. The conditional deletion of PfCDP leads to the selective up-regulation of a subset of *rifins* in *P. falciparum*. Further, *in vitro* rosetting assay has revealed that PfCDP null *P. falciparum* lines promote rosette formation in RBCs. Together, this study reports on the epigenetic regulator that controls the expression of subsets of important virulent genes in *P. falciparum* and provides evidence on the molecular basis of the subsets of RIFIN expression and RBC rosetting.

## Materials & Methods

### *P. falciparum* culture maintenance and transfection

The *P. falciparum* parasite lines used in this study were derived from the NF54 strain. Asexual blood stages of *P. falciparum* were cultured at 5% haematocrit using advanced RPMI 1640 (Gibco cat # 12633012) supplemented with 2 mM L- glutamine (Gibco # 2530081) 10% of O+ve plasma, 0.6% Albumax II (Gibco # 11021-037), 25 mM HEPES (Millipore # 391338), and 100 µM hypoxanthine (Sigma-Aldrich # H9636). The Parasites were cultured in O^+ve^ red blood cells incubated at 37 °C, under a gas phase of 5% CO_2._ The collected parasites were released from infected RBCs by lysis with 0.15 % saponin (Sigma-Aldrich # 47036). The lysed RBCs were pelleted at 14,000 RPM for 10 min at 4 °C to separate the parasites and further washed with ice-cold (Phosphate buffered saline) PBS 3 times and stored at -80 °C for further studies. The RPMI 1640 medium was pre-warmed to 37°C before usage, and media change was done every alternate day. The parasite growth was regularly monitored by microscopy using standard Giemsa-stained blood smears. The parasite synchronization was done by Percoll enrichment. For transfection, we used the Amaxa™ 4D-Nucleofector™ technology with purified NF54-DiCre schizonts, 50-60 μg of linearized repair plasmid, and 25-30 μg of the CRISPR/Cas9 plasmid containing the guide RNA. Transfection of linear DNA has the advantage of not retaining the linear DNA beyond 4 d post-transfection (33). For this reason and the marker-free approach chosen here, hDHFR selection with 5 nM WR99210 (Jacobus Pharmaceuticals) was applied for 4 d. Viable parasites were detected 15 d post-transfection and subsequently screened for recodonized chromodomain (CD) integration by PCR.

### *P. falciparum* recombinant chromodomain expression and purification

The chromodomain (amino acids 42 – 108) present in the conserved chromoprotein (PF3D7_1140700) was amplified using domain-specific primers (Supplementary table 2). The amplicons were then cloned into the pGEX6P2 vector (GE Healthcare Life Sciences) using BamHI and XhoI sites. All clones were sequenced to confirm the frame alignment. The expression vector pGEX6P2 – PfCD was transformed into host *E. coli* BL21 cells. A single colony was inoculated into Luria-Bertani broth containing ampicillin and incubated at 37 °C with constant agitation (200 rpm). The next day 1% of the overnight grown culture was inoculated to 1 litre of LB medium containing the ampicillin. The cells were grown in LB medium until the mid-exponential phase (OD_600_ 0.5–0.6), and the expression was induced by the addition of 1 mM isopropyl-β-D-1-thiogalactopyranoside (IPTG) (Sigma-Aldrich # I6758). After the addition of IPTG, the culture was shifted to 18 °C and incubated overnight with shaking. The following day cells were pelleted at 4000 rpm, 4 °C for 45 min. The pellet was washed in a buffer containing 20 mM Tris-HCl pH 8.0, 50 mM NaCl and water and stored at -20 °C. The protein was expressed as a fusion protein containing an amino-terminal GST-tag, and soluble GST fusion proteins were purified easily using an immobilized glutathione Sepharose (GE # 17-5279-01) column. The collected cell pellet was resuspended in sonication buffer (20 mM HEPES, 500 mM KCl, 1 mM EDTA, 1 mM DTT, 10% Glycerol) PMSF, bacterial protease inhibitor cocktail and thoroughly mixed, and lysed with a sonicator (Sonics Vibra cell) using 38% amplitude with pulse 1 s on and 1 s off. After sonication, the soluble fraction was collected by high-speed centrifugation at 12000 rpm at 4°C for an hour and passed onto the column packed with Glutathione Sepharose^TM^ High-performance column. The column was washed with a sonication buffer, and the bound proteins were eluted using a sonication buffer containing 40 mM reduced Glutathione. The proteins were dialyzed against buffer I (20 mM HEPES pH 7.5, 200 mM KCl, 1 mM DTT, 1 mM EDTA, and 10% glycerol) for 2 h at 4 °C and subsequently dialyzed against buffer II (20 mM HEPES pH 7.5, 200 mM KCl, 1 mM EDTA, 1 mM DTT and 60% glycerol). The protein concentration was measured using nanodrop, and purity was analyzed on an SDS-PAGE gel, stained with Coomassie brilliant blue stain. The purified proteins were confirmed by western blotting using an anti-GST antibody (Cell Signaling Technology # 91G1).

### Site directed mutagenesis, expression and purification of mutant PfCD domains

We performed site-directed mutagenesis with the pGEX6P2-PfCD clone was performed to introduce alanine at selected aromatic amino acid residues (Supplementary table 2) to generate pGEX6P2-PfCD F75A, pGEX6P2-PfCD W82A, pGEX6P2-PfCD Y89A mutant clones. The mutants were generated with primers containing mutation by using the megaprimers followed by rolling circle amplification PCR. The presence of appropriate mutations was confirmed by Sanger sequencing. The mutant domain clones were expressed in *E. coli* BL21 strain and purified using Glutathione SepharoseTM High-performance column as described above. The quality of the mutant proteins was analyzed on 12% SDS-PAGE gel and stained with Coomassie brilliant blue. The homology model for PfCD domain protein was generated with human HP1 chromodomain as a template using Swiss-Model with default parameters. Docking of H3K9me3 peptide with PfCD structure model was developed using pyDockWEB (https://life.bsc.es/pid/pydockweb) with default parameters. The modelled PfCD – H3K9me3 peptide structure was visualized using pyMOL software.

### Screening for methylation interaction using Modified Histone Peptide Array

MODified Histone peptide arrays (Active Motif™) are used to screen the protein and histone modification interactions. The peptide Array contains 384 modifications (acetylation, methylation, phosphorylation, and citrullination) on the N-terminal tails of histones H2A, H2B, H3, and H4. The array was blocked overnight at 4 °C in 5% non-fat dried milk in Tris buffer saline containing 0.05% Tween 20, and 150 mM NaCl. The array was washed in TTBS (Tris Buffered saline containing 0.05% tween) buffer three times for 5 min at room temperature, and 1 μM of GST tagged PfCD protein was added to the array in interaction buffer (100 mM KCl, 20 mM HEPES PH 7.5, 1 mM EDTA, 0.1 mM DTT and 10% Glycerol), and incubated for 2 h. After the incubation, the array was washed in Tris-buffered saline containing 0.05% tween three times. Slides were first incubated with anti-GST primary antibody (1:5000) for 1 h at room temperature and subsequently probed with appropriate secondary antibodies coupled to Horseradish peroxidase for 1 h at room temperature followed by chemiluminescent detection with ECL reagent (Bio-Rad).

### Peptide pull-down assay

Histone peptides can be used to identify the potential readers of histone modifications very specifically in an unbiased fashion. Biotinylated histone peptides that are either unmodified or modified at specific residues are immobilized on avidin beads and incubated with recombinant proteins (Supplementary table 3). We used biotinylated histone peptides for the pull-down assays, unmodified histone H3, histone H3K9me3 (Anaspec) were used for the experiment. These Peptides (35 µM) were incubated with Dynabeads™ M-280 streptavidin (Invitrogen # 11205D) in binding buffer (20 mM HEPES pH 7.5, 10% v/v glycerol, 100 mM KCl, 5 mM MgCl2, 1 mM EDTA and 1 mM DTT) and incubated in a roto spin to mix briefly, followed by washing with the same buffer for 4 times. Then PfCD and mutant proteins (1 µM) were incubated with the beads for 3 h at room temperature with proper mixing, further washed 5 times with wash buffer (20 mM HEPES pH 7.5, 10% v/v glycerol, 200 mM KCl, 5 mM MgCl2, 1 mM EDTA and 1 mM DTT) containing high salt concentration to remove the unbound and non-specific binding proteins. The beads were resuspended in 1X PBS containing Laemmli buffer and boiled for 15 min at 95 °C. The bound proteins were analyzed by resolving it on 12% SDS-PAGE and stained with Coomassie brilliant blue.

### *P. falciparum* mononucleosome pull down assay

Mononucleosomes were prepared from *P. falciparum*, and the quality was analyzed on 1.5% Agarose gel (Supplementary figure 3A). We used these mononucleosomes were used for the nucleosome pull-down studies using GST pull-down assays for PfCD wild type and mutant proteins. In nucleosome pull-down assay, we used 1 µM PfCD wild type and the mutant proteins and incubated with isolated mononucleosomes in interaction buffer (20 mM HEPES pH 7.5, 10% v/v glycerol, 100 mM KCl, 5 mM MgCl2, 1 mM EDTA, and 1mM DTT) and incubated at 4 °C for 4 h. For the control assays, we used recombinant GST protein that was incubated with nucleosomes. The PfCD – *P. falciparum* nucleosomes complex was further incubated with glutathione Sepharose beads for 2 h using interaction buffer. Following incubation, the beads were washed thoroughly with wash buffer (20 mM HEPES pH 7.5, 10% v/v glycerol, 200 mM KCl, 5 mM MgCl2, 1 mM EDTA, and 1 mM DTT) to remove the unbound protein complexes. The bound protein and nucleosome complexes were separated on 16% SDS-PAGE gel and subjected to western blot analysis. The presence of various histone methylation marks with PfCD bound nucleosome complexes were analyzed using modification specific antibodies against H3K9me3 (Diagenode # C15410193), H3K36me3 (Diagenode # C15410192), and H3K4me3 (Diagenode # C15410003). The blot was cut above 22 kDa and probed with an anti-GST antibody to confirm an equal concentration of the GST proteins in the complexes.

### Peptide pull-down and dot blot assay

Biotinylated unmodified histone H3 and biotinylated histone H3K9me3 peptides (10 µM) were incubated with Dynabeads™ M-280 streptavidin for 1 h in an interaction buffer with 20 rpm. After washing four times with wash buffer, the complex was allowed to interact with varying PfCD protein concentrations (25 nM-300 nM) for 1 h. The complex was washed extensively with wash buffer, and the beads were resuspended in 1X PBS containing Laemmli buffer and boiled for 15 min at 95 °C. After the pull-down, the samples were spotted on a nitrocellulose membrane (Cat # 1620115), carefully using a narrow-mouth pipette tip. The membrane was allowed to dry and the samples were cross-linked to a nitrocellulose membrane using a UV cross linker UVI Link CL-508 (0.1 J for 6 min). The membrane was further blocked with 5% skimmed milk for 2 hours, followed by TTBS wash three times for 5 min at room temperature. The membrane was probed with anti-GST primary antibody (1:6000) for 2 h and subsequently probed with appropriate secondary antibody coupled to Horseradish peroxidase for 2 h at room temperature followed by chemiluminescent detection with ECL reagent.

### Mouse immunization and serum preparation for PfCD domain

We used the C57/BL6 mouse stain to generate polyclonal antibodies for the PfCD domain. The primary immunization was carried out using 450 μg of GST cleaved PfCD protein and an equal volume of Freund’s complete adjuvant (F5881 # Sigma-Aldrich) via the intraperitoneal route. The second, third, and fourth booster immunizations were done via the intraperitoneal route using Freund’s incomplete adjuvant (F5506 # Sigma-Aldrich). The mouse was sacrificed, and the blood was collected and kept at room temperature for 2 h, followed by incubation at 4 °C overnight to collect the serum. The next day the sample was centrifuged at 2000 rpm for 10 min at 4 °C to remove the contaminant RBCs; the serum was obtained in the supernatant and again centrifuged at 14000 rpm for 15 min to obtain clear serum and stored at -80 °C. All animal procedures were performed according to the guidelines approved by the RGCB institutional ethics committee.

### Purification of PfCD IgG from immune serum

We used Dynabeads® Protein A for the purification of mouse IgG from the immunized serum. The beads were washed thoroughly with 1X PBS and pre equilibrated with a wash buffer (100 mM Tris and 135 mM NaCl). The serum was mixed in 1X PBS and incubated with Dynabeads® Protein A (Invitrogen # 10001D) at 4 °C for 5 h. After the incubation, the supernatant was discarded and washed with a wash buffer (100 mM Tris and 135 mM NaCl). The elution of bound IgGs was performed using the elution buffer (100 mM glycine pH 3.0) and quickly transferred to a fresh Eppendorf tube containing 2 µl of neutralization buffer (2 M Tris pH 9) with efficient mixing. The antibody was aliquoted and stored at -80 °C for further use. The collected anti-PfCD IgG fractions were resolved by 12% SDS-PAGE and stained using Coomassie brilliant blue.

### Chromatin Immunoprecipitation (ChIP) assay

The chromatin was prepared from the *P. falciparum* trophozoite stage; the cells were cross-linked using 1% formaldehyde for 10 min, followed by the addition of 125 mM of glycine and incubated for 5 min. The nuclei were isolated from fixed *P. falciparum* cells by homogenization in buffer 20 mM Tris pH 8.0, 6 mM MgCl_2_ and 0.4% Nonidet P-40, and collected on a 0.5 M sucrose-buffer cushion and suspended in SDS buffer (2% SDS, 100 mM Tris, pH 8.0, 20 mM EDTA, protease inhibitors). The chromatin was sheared by sonication in a Bioruptor UCD-200 (Diagenode) for 10 min, at 30 s intervals, and the quality of the chromatin was analyzed on 1.5% agarose gel to get the 200 - 500 bp. We performed ChIP using an anti-PfCD IgG or control mouse pre-immunesera were added to the ChIP buffer (100 mM NaCl, 20 mM Tris pH-7.5, 6 mM EDTA, 1% Triton X-100) containing cross-linked chromatin samples and, incubated at 4 °C for overnight. The Protein A Dynabeads (15 μl) were blocked using 3% BSA for 1 h at 4 °C and, the beads were added to the samples, further incubated for 2 h at 4 °C with proper mixing. The bound complexes were washed with a wash buffer (180 mM NaCl, 20 mM Tris pH-7.5, 6 mM EDTA, 1% Triton X 100) four times to remove the unbound proteins. The ChIP complexes were diluted in sterile 1X PBS and boiled at 95 °C with Laemmli buffer. The samples were resolved on 16% SDS-PAGE, followed by western blot analysis. The Presence of various methylation marks in the ChIP complexes was analyzed using antibodies against H3K9me3, H3K36me3, H3K4me3 marks, and the anti-histone H3 antibody (Diagenode # C15210011) was used as the loading control.

### Generation of CRIPSR Cas9 PfCD gRNA constructs for conditional gene deletion

We have used a recently developed CRISPR cas9 system that allows the conditional, and site-specific elimination of genomic sequences of essential and nonessential genes in *P. falciparum* (Primers used to generate the KO constructs are provided in Supplementary table 2). This is achieved by integrating loxP sites into a short synthetic intron to create a unit (loxPint), incorporated anywhere in open reading frames without hampering the protein expression. This, along with the Rapamycin-induced activation of dimerizable Cre recombinase, forms a more reliable gene inactivation system (34) (35). We used two different plasmids for this; One is a pUC-18 plasmid carrying a recodonized PfCD donor DNA segment flanked by loxPint sites and a pDC_Cas9_hDHFRyFCU. The recodonization of PfCD was performed to maintain the codons for the same amino acids but different DNA sequence, which permits to screening the clones after the integration into the genome of *P. falciparum*. The chromodomain region was recodonized, and the loxPint sites were incorporated to the 5’ and 3’ ends, each flanked by the homologous regions for 5’ and 3’ regions that further assist the process of homologous recombination. The recodonized chromodomain was inserted into the pUC18 vector using BamHI and HindIII restriction sites (Supplementary figures 4A – 4B). The guide RNA was designed for the wild type chromodomain and cloned into the pDC_Cas9_hDHFRyFCU using the BbsI enzyme, as previously described (36). Both constructs were sequenced and confirmed (Supplementary figures 4C – 4D). To induce DiCre-driven loxP site recombination, parasites were treated with 100 nM Rapamycin (Sigma-Aldrich # R8781) or dimethyl sulfoxide (DMSO) (Invitrogen # D12345) (0.1% [vol/vol]) for 4 h. The Parasites were subsequently washed twice with warm RPMI 1640 medium and returned to culture. Samples used for nucleic acid extraction and protein extraction were taken at least 48 h after Rapamycin or DMSO treatment. The deletion of the PfCD fragment was confirmed by PCR and the loss of protein was confirmed by immunoblot using anti-PfCD antibody, the anti-Plasmodium aldolase antibody (abcam # ab207494) was used as the loading control.

#### Sample preparation for transcriptome analysis for wild type and PfΔCDP *P. falciparum*

The PfΔCDP *P. falciparum* parasite lines were grown at 5% haematocrit in complete cell culture media, as described earlier. The conditional deletion of the PfCDP was performed by 100 nM Rapamycin treatment, while the control cells were treated with DMSO. The parasites were released from infected RBCs 0.15% saponin lysis, and RNA was isolated from the control and PfΔCDP *P. falciparum* with Trizol solution followed by Qiagen RNA isolation kit (Cat # 74104). The quality of the RNA quality was analyzed on the formaldehyde-denatured gel, and the RNA was stored at − 80 °C.

### RNA Quality Control and library preparation

The RNA quantity and quality was assessed using Nanodrop2000 (Thermo Scientific, USA), Qubit (Thermo Scientific, USA) and Bioanalyzer 2100 (Agilent, USA). The RNA sequencing libraries were prepared with Illumina-compatible NEBNext® Ultra™ II Directional RNA Library Prep Kit (New England BioLabs, MA, USA) at Genotypic Technology Pvt. Ltd., Bangalore, India. Briefly, 100 ng -500 ng of Qubit quantified total RNA was taken for mRNA isolation, fragmentation and priming. Fragmented and primed mRNA was further subjected to first -strand cDNA synthesis followed by second -strand synthesis. The double -stranded cDNA was purified using JetSeq Beads (Bioline, Cat # BIO-68031). The purified cDNA was end-repaired, adenylated, and ligated to Illumina multiplex barcode adapters as per NEBNext® Ultra™ II Directional RNA Library prep protocol followed by the second -strand excision using USER enzyme at 37 °C for 15 min.

#### Illumina Universal Adapters

5’AATGATACGGCGACCACCGAGATCTACACTCTTTCCCTACACGACGCTCTTCCG ATCT-3’

#### Index Adapter

5’-GATCGGAAGAGCACACGTCTGAACTCCAGTCAC-[INDEX]-ATCTCGTATGCCGTCTTCTGCTTG-3’.

[INDEX] –Unique sequence to identify sample-specific sequencing data.

Adapter ligated cDNA was purified using JetSeq Beads and was subjected to 11-12 cycles for Indexing (98 °C for 30 s, cycling (98 °C for 10 s, 65 °C for 75 s) and 65 °C for 5 min to enrich the adapter-ligated fragments. The final PCR product (sequencing library) was purified with JetSeq Beads (Bioline, Cat # BIO-68031), followed by library quality control check. Illumina-compatible sequencing libraries were quantified by Qubit fluorometer (Thermo Fisher Scientific, MA, USA) and its fragment size distribution was analyzed on Agilent 2200 TapeStation.

### Illumina Sequencing

The libraries were sequenced on Illumina HiSeq 4000 sequencer (Illumina, San Diego, USA) for 150 bp paired-end chemistry following the manufacturer’s procedure. The data obtained from the sequencing run was de-multiplexed using Bcl2fastq software v2.20 and FastQ files were generated based on the unique dual barcode sequences.

### Quality Check and trimming of the sequencing reads

The sequencing quality was assessed using the FastQC software ver. 0.11.9. The adaptor sequences and low-quality bases (Phred quality cut off set at 30) were trimmed from the reads using the Trim Galore tool ver. 0.6.5.

### Alignment of the reads to the reference genome

The reads that survived quality checks were aligned to the *Plasmodium falciparum* 3D7 isolate genome (ver. 46) using the Hisat2 read alignment tool. The library strandedness was set as the first-strand in accordance with the strand-specific library preparation protocol. The alignment output files in the sam format were sorted and converted to the bam format using the Samtools utilities (ver 1.10).

### Quantification of transcript abundance and differential gene expression analysis

The transcript abundance count data was generated using the HTSeq-count function (ver. 0.12.4). The differential gene expression analysis was performed using the DESeq2 package (ver. 3.11) in R programming environment. Genes were classified as deregulated based on log_2_fold-change cut off of +1 (for up-regulation) and -1 (for down regulation) and a p-value cut off of 0.05.The Gene Ontology analysis was performed using the GO tool hosted by the PlasmoDB web database and data visualization of the DGE was performed using custom scripts in the R programming environment and with DESeq2 package utilities.

### ChIP-sequencing for identification of the occupancy of PfCDP on *P. falciparum* genome

For ChIP sequencing analysis, we used 12 µg of crosslinked *P. falciparum* chromatin and 2 µg control IgG, anti-PfCD IgG or control mouse pre-immune sera to perform the immunoprecipitation experiment. The anti-PfCD IgG and *P. falciparum* chromatin were allowed to interact overnight at 4 °C in ChIP buffer. To the sample, 15 µl of protein A beads were added and incubated for 2 h at 4 °C. The complex was washed extensively using wash buffer and the immuno-precipitated DNA was eluted and purified using PCR purification columns (MN-740609) and quantified. The ChIP libraries were prepared using NEBNext® UltraTM II DNA Library Prep Kit for Illumina (Catalog: E7645S, New England Biolabs). The samples were subjected to various enzymatic steps to repair the ends and tailing with dA-tail followed by an adapter ligation. To the adenylated fragments, loop adapters were ligated and cleaved with the uracil-specific excision reagent (USER) enzyme. The samples were further purified using 0.9X AMPure XP beads (Catalog: A63881, Beckman Coulter). Furthermore, the DNA was amplified by 12 cycles of PCR with the addition of NEBNext Ultra II Q5 master mix, and “NEBNext® Multiplex Oligos for Illumina” to facilitate multiplexing while sequencing. The amplified products were then purified using 0.9X AMPure XP beads (Catalog: A63881, Beckman Coulter) and the final DNA library was eluted in 15μl of 0.1X TE buffer. The library quality assessment was done using Agilent D1000 Screen Tape System (Catalog: 5067-5582, Agilent) in a 4150 Tape Station System (Catalog: G2992AA, Agilent) and subjected to paired-end sequencing using Illumina HiSeq platform. The ChIP sequencing and analysis for PfCDP was performed in duplicates, and we have used pooled input samples for the normalization.

### ChIP sequencing analysis

The ChIP and input seq reads were quality checked using FASTQC and trimmed to remove subpar sequences and adapters using Trim_galore! Alignment to the *P. falciparum* reference genome was performed using Bowtie2 and the aligned reads were sorted using SAMTools. Peak calling was performed using the MACS3 software using the peakcall option in no-model mode. The FDR/q-value was used 5.00e-02 for scoring the peaks. Peaks with greater than 1.8 fold enrichment over input were taken for consideration with high confidence attributed to those with 2 fold enrichment and above. The H3K9me3 ChIP-seq dataset was obtained from the Gene Expression Omnibus database. (https://www.ncbi.nlm.nih.gov/geo/query/acc.cgi?acc=GSE63369). Gene Ontology was performed using the online tool available on PlasmoDB web resource (https://plasmodb.org/plasmo/app). Deeptools was used for visualization and additional data analysis. Bamcoverage tool was used to generate bigwig files from the normalized ChIP datasets. The computeMatrix option was used to generate the average gene occupancy profile over the PfCDP1 target genes and their flanking regions. The matrix was imputed into the plotHeatmap option to generate the PfCDP1 occupancy heatmap. The compute Matrix tool was further also used to generate a comparative gene occupancy profile for PfCDP1 and the H3K9me3 histone modification (using virulence and non-virulence gene bed files as reference) and visualized using the plotProfile option. The PfCDP1 and H3K9me3 bigwig files were compared by the multiBigwigSummary tool to evaluate correlation between their pan genome occupancy across defined bins. The PfCDP1 genomic occupancy dataset and gene expression dataset from wildtype *P. falciparum* lines were compared using the multiBigwigSummary tool to assess the correlation of PfCDP1 genomic occupancy with target gene expression. R Studio was used to generate boxplots and scatterplots representations of these aforementioned analyses.

### Rosette formation assay for PfΔCDP and wild type *P. falciparum*

The PfΔCDP *P. falciparum* parasite lines were cultured in fresh human O+ve RBC at 5% haematocrit in a complete cell culture medium (CCM – RPMI medium with Albumax II). The conditional deletion of Pf CDP was performed by treating the cells with 100 nM Rapamycin, while the control cells were treated with the same concentration of DMSO. Media change was done every day without disturbing the sedimented erythrocytes. The rosette forming infected RBCs (iRBC) were enriched from control and PfΔCDP *P. falciparum* culture as described (37). Briefly, we harvested the *P. falciparum* culture containing mature trophozoite and schizont stages, having 5-8% parasitaemia by centrifuging the culture for 10 min at 1,200 rpm using a swing bucket rotor. After discarding the supernatant, the pellet was resuspended carefully in fresh CCM (We used 5.5 ml CCM to dissolve 0.5 ml of the pellet). The resuspended culture (2 ml each) was carefully overlaid on top of 2 ml Histopaque®-1077 and centrifuged immediately for 30 s at 2400 rpm at room temperature. After centrifugation, the supernatant was removed, and the pellets were pooled and resuspended in 2 ml of CCM. For the assessment of the rosetting rate, we stained the *P. falciparum* using Hoechst 33342. A 1.5 ml round bottom tube 100 μl of CCM containing 10 μg/ml Hoechst was taken and added 2-3 μl of prepared cell pellet. The tube was gently mixed and incubated in the dark for 10-12 min for nuclear staining, then the pellet was washed by the addition of 900 μl CCM, followed by centrifugation at 1200 rpm for 2 min at room temperature. The supernatant was removed very carefully, and the pellet was resuspended in 40 μl of CCM. A clean microscope glass slide was taken and carefully placed 10 μl rosette suspension onto it and covered it with a clean coverslip. We sealed the coverslip’s edges by using nail polish and examined the slides under a fluorescence microscope using a 60X objective and used both UV and bright-field light to visualize both iRBCs and RBC. We Counted 230-250 parasite iRBCs and scored the number of iRBCs engaged in rosettes (mature stages having bound two or more uninfected RBC). We calculated the percentage of the rosetting using the formula, Rosetting rate = (Number mature stage iRBC engaged in rosettes/Number of mature stages) X 100 (37).

## Results

### Identification and characterization of H3K9me3 specific methyl reader protein from *P. falciparum*

The H3K9me3 is an important repressor methyl mark for virulence family genes in *P. falciparum*; to date, only reader protein for this mark, HP1 chromodomain was characterized (24, 28, 38). Genetic knockout of HP1 chromodomain in *P. falciparum* does not lead to the up-regulation of all virulence family genes, suggesting the presence of other essential H3K9me3 methyl-binding proteins in *P. falciparum*. To identify other potential H3K9me3 binding proteins, we performed sequence analysis of *P. falciparum* chromodomain with the well-characterized H3K9me3 methyl reader proteins of other eukaryotes. The chromodomain of *P. falciparum* is conserved for the aromatic amino acids that form a methyl binding pocket (Supplementary figure 1A). We cloned, expressed and purified the uncharacterized putative chromodomain (CD) of chromodomain-containing protein (PfCDP) (PF3D7_1140700) from *P. falciparum* (Figure 1A; Supplementary figure 1B). The binding specificity of the recombinant PfCD protein was analysed by using a MODified histone peptide array (Active Motif, Belgium), which contains 384 histone modifications and permits screening the histone modifications specific interaction of methyl-reader proteins under a competitive environment. The modified histone peptide array contains all physiologically relevant modifications, the modified histone tails are unstructured thus mimicking the cellular conditions (39). A far-western blot approach was used to study the interaction of purified GST tagged recombinant chromodomain (PfCD) on the peptide array. The PfCD selectively interacted with the H3K9me3, H3K27me2, and H3K27me3 marks containing peptides (Figure 1B). In addition, we found interaction of PfCD with peptides containing H3R2me2a, H3R8me2s modifications along with the H3K9me3 mark (Figure 1B; Supplementary figure 2). We also observed interactions of the PfCD with H4 acetylation modification peptides on the array (Supplementary figure 2). In addition, we observed interaction with the H3K27me2 and H3K27me3 marks on the peptide array, however these methyl marks are not well characterized in *P. falciparum* (7, 40) The commonly observed modification in all the intense spots on the array is H3K9me3, therefore, we focused only on PfCD – H3K9me3 mark interaction. To confirm the H3K9me3 interaction by PfCD, we performed a peptide pull-down assay using a synthetic peptide containing H3K9me3 modification and an unmodified H3 peptide as a control. We observed considerably better binding of PfCD to the H3K9me3 peptide than to the unmodified H3 peptide (Figure 1C). To further substantiate the H3K9me3-PfCD interaction, we performed homology modeling with known characterized eukaryotic chromodomain proteins and followed by docking studies with the H3K9me3 peptide to identify the methyl-binding pocket, with characteristic aromatic amino acids in PfCD (Supplementary figure 1C) (41, 42). Based on the model, we mutated the aromatic amino acid residues F75A, Y82A, W89A, and purified the recombinant PfCD mutants as GST tagged protein with equal quality to that of the wild type protein (Supplementary figure 1D). Peptide-pull-down assays were performed with these purified recombinant PfCD mutant proteins along with the wild type protein. The PfCD wild type protein retained binding to the H3K9me3 peptide, whereas the three mutant proteins showed a considerable reduction in binding (Figure 1D). Since the PfCD protein also binds to unmodified H3K9 peptides, but less so, we measured the binding affinity of PfCD to the H3K9me0 and H3K9me3 peptides. We performed the peptide pull-down assay with varying concentrations of the PfCD protein, and the bound fractions were determined by dot blot assay. The assay has identified that PfCD binds to H3K9me3 peptide with more than 3-fold higher affinity than to unmodified H3K9me0 peptide (Figures 1E – 1G). There were significant differences with PfCD protein bound fractions between modified and unmodified peptide and confirms that the PfCD binds to H3K9me3 peptide with higher affinity (Figures 1E–1G). Next, we studied the interaction of PfCD with mononucleosome preparations from the trophozoite stages of *P. falciparum* (Supplementary figure 3A). The *in vitro* mononucleosome binding assay suggests that PfCD binds strongly to the H3K9me3 mark containing nucleosomes and does not bind to nucleosome containing an active methylation mark. We also observed a weak binding to H3K36me3 containing nucleosomes; this could be due to the co-existence of both marks on a single nucleosome (Figures 2A–2B). We verified the nucleosome interaction with the PfCD mutant proteins and found a significant reduction in mutant protein binding with H3K9me3 containing nucleosomes (Figure 2C–2D). Taken together, through *in vitro* binding assays, we have confirmed that the PfCD can interact with the H3K9me3 mark. Such modification specific interactions may regulate the complex processes of gene expression in *P. falciparum*.

**Figure 1:**
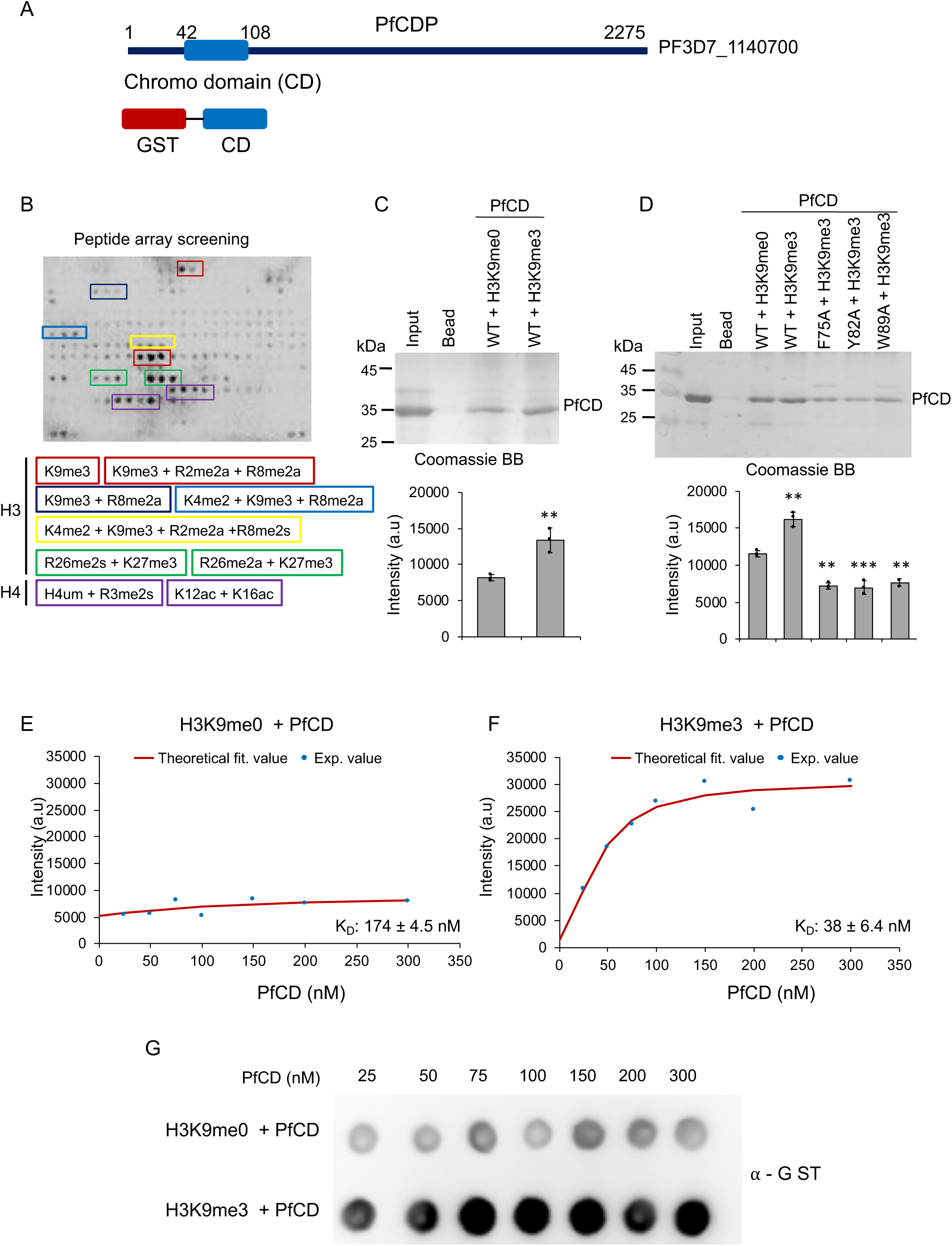
Screening, identification and validation of H3K9me3 methyl-binding protein from *P. falciparum*. **(A)** Schematic representation of PfCDP, the chromodomain (CD) marked with a blue color box. The bottom scheme represents the GST tagged PfCD protein used for all peptide array and binding assays. **(B)** Peptide array screening with GST – PfCD recombinant protein (1 μM) and interaction was confirmed using an anti-GST antibody. The red boxes represent the H3K9me3 containing spots, the blue color box represents the H3K9me3 and H3R8me2 peptides, green color boxes represent H3R26 and H3K27 methylated peptides. **(C)** Peptide pull-down assay with biotinylated H3K9me0 and H3K9me3 peptides as bait and GST-PfCD wild type protein as prey. The peptide bound protein complexes were separated on 12% SDS-PAGE gel and stained with Coomassie brilliant blue. The bottom bar plot represents the band intensity analyzed using ImageJ, and the error bar represents the SEM of three independent pull -down assays. The p values are calculated using paired t-test. **(D)** Peptide pull-down assay with biotinylated H3K9me3 peptides as bait and GST-PfCD wild type and methyl-binding pocket mutant proteins as prey. The peptide-bound protein complexes were separated on 12% SDS-PAGE gel and stained with Coomassie brilliant blue. The bottom bar plot represents the band intensity analyzed using ImageJ, and the error bar represents the SEM of three independent pull-down assays. The p values are calculated using paired t-test. **(E)** The peptide pull-down assay with H3K9me0 peptide was performed with increasing concentration of PfCD protein. **(F)** The peptide pull-down assay with the H3K9me3 peptide with an increasing concentration of PfCD protein. **(G)** A representative dot blot image from a single pull down assay and the peptide bound fractions were measured by dot blot assay and probed with an anti-GST antibody. The peptide-down assay was performed in triplicate and the intensity of each spots for all the experiments was measured by ImageJ and the values were plotted in excel to determine the equilibrium binding constant using the Microsoft excel solver module. The K_D_ values are provided in the each graph (PHD + H3K9me0 is 175 nM; PHD + H3K9me3 is 38 nM) and the experimental deviation in the dissociation constant is provided in the graph.

**Figure 2:**
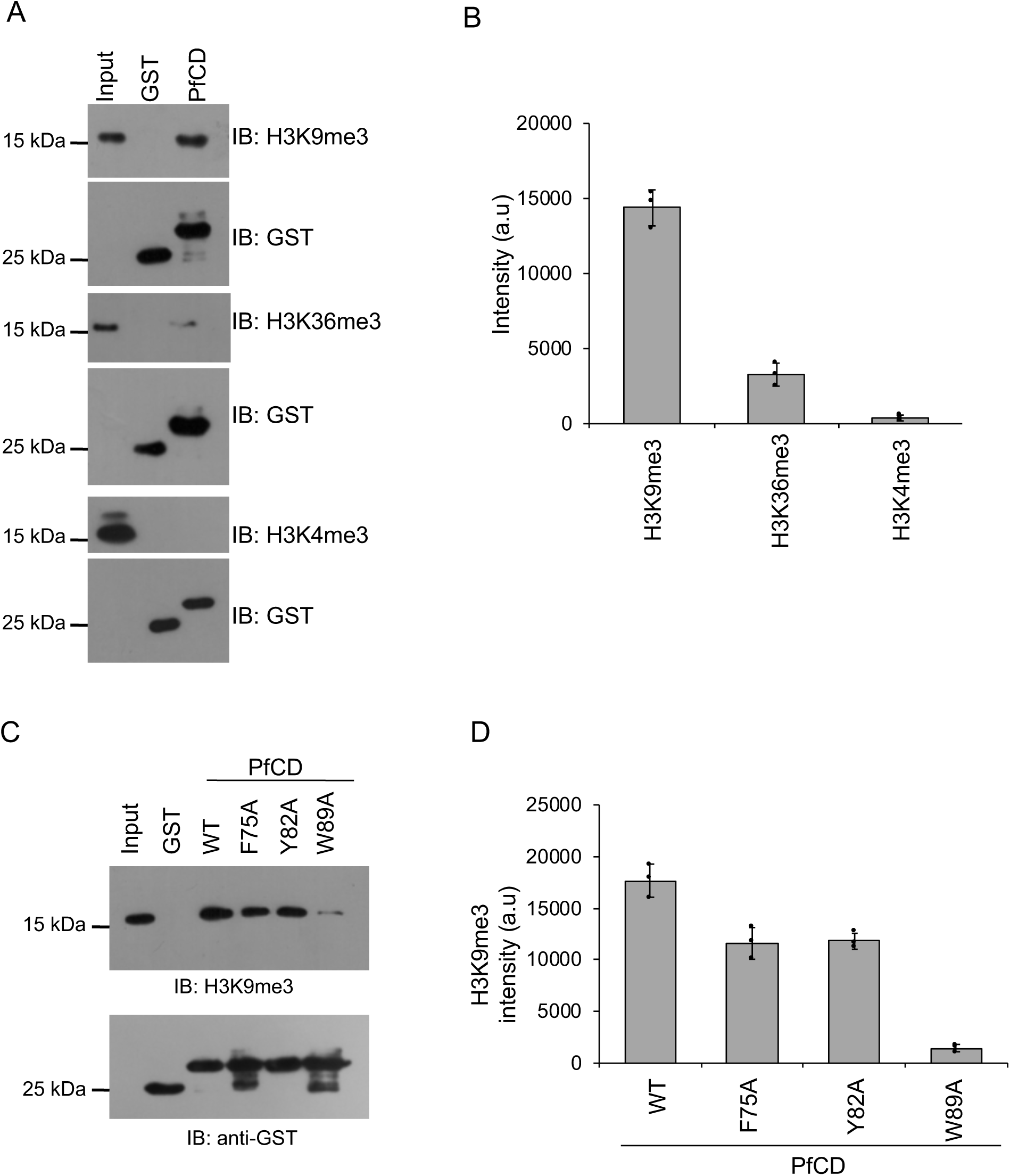
The PfCD protein binds to H3K9me3 containing nucleosome substrates. **(A)** The GST-PfCD pull-down assay was performed using mononucleosomes isolated from *P. falciparum,* and protein-bound nucleosomes complexes were separated on 16% SDS-PAGE gel, transferred to a PVDF membrane, probed with anti-H3K9me3, anti-H3K36me3 and anti-H3K4me3 antibodies. The bottom blot represents the immunoblot of GST and GST-PfCD proteins as loading controls from the same experiment. **(B)** The bar plot represents the band intensity of H3K9me3, H3K36me3 and, H3K4me3 methyl marks which were measured using ImageJ, and the error bar represents the SEM of three independent pull-down assays. **(C)** The GST-PfCD wild type and mutant proteins pull-down assay with mononucleosomes and protein-bound nucleosomes were separated on 16% SDS-PAGE gel, transferred to a membrane, probed with the anti-H3K9me3 antibody. The Bottom western blot image represents the immunoblot of GST and GST-PfCD proteins using an anti-GST antibody as loading controls from the same experiment. (**D**) The bar plot represents the band intensity of H3K9me3 methyl mark, which was measured using ImageJ, and the error bar represents the SEM of three independent pull-down assays.

### PfCDP interacts to the H3K9me3 mark on chromatin of *P. falciparum*

We next tested whether PfCDP binds to H3K9me3 modifications on chromatin in *P. falciparum* infected red blood cells. To do that, we generated a chromodomain (CD) specific antibody in mice using recombinant PfCD protein (Supplementary figure 3B). The western blot assay using PfCD recombinant protein with purified anti-PfCD IgG (subsequently called anti-PfCD) confirms its binding. No signal was observed with pre-immune sera (Supplementary figure 3C). Purified anti-PfCD binds to endogenous PfCDP in *P. falciparum* lysate showing a band of the expected size of PfCDP of 250 kDa, whereas no signal was observed with pre-immune sera (Figure 3A). To study the endogenous interaction of PfCDP and the H3K9me3 mark on the chromatin of *P. falciparum*, we performed chromatin immunoprecipitation (ChIP) using anti-PfCD antibody and pre-immune sera as control (Supplementary figure 3D). The immunoblot with anti-H3K9me3 antibody confirms that the PfCDP co-precipitates H3K9me3 containing chromatin from *P. falciparum* (Figure 3B), and does not co-precipitate the well-known gene activation methyl mark H3K4me3 mark containing chromatin (Figure 3B – 3C). The cellular ChIP assay establish that the PfCDP binds to the H3K9me3 mark containing chromatin in *P. falciparum,* indicating its possible role in gene regulation.

**Figure 3:**
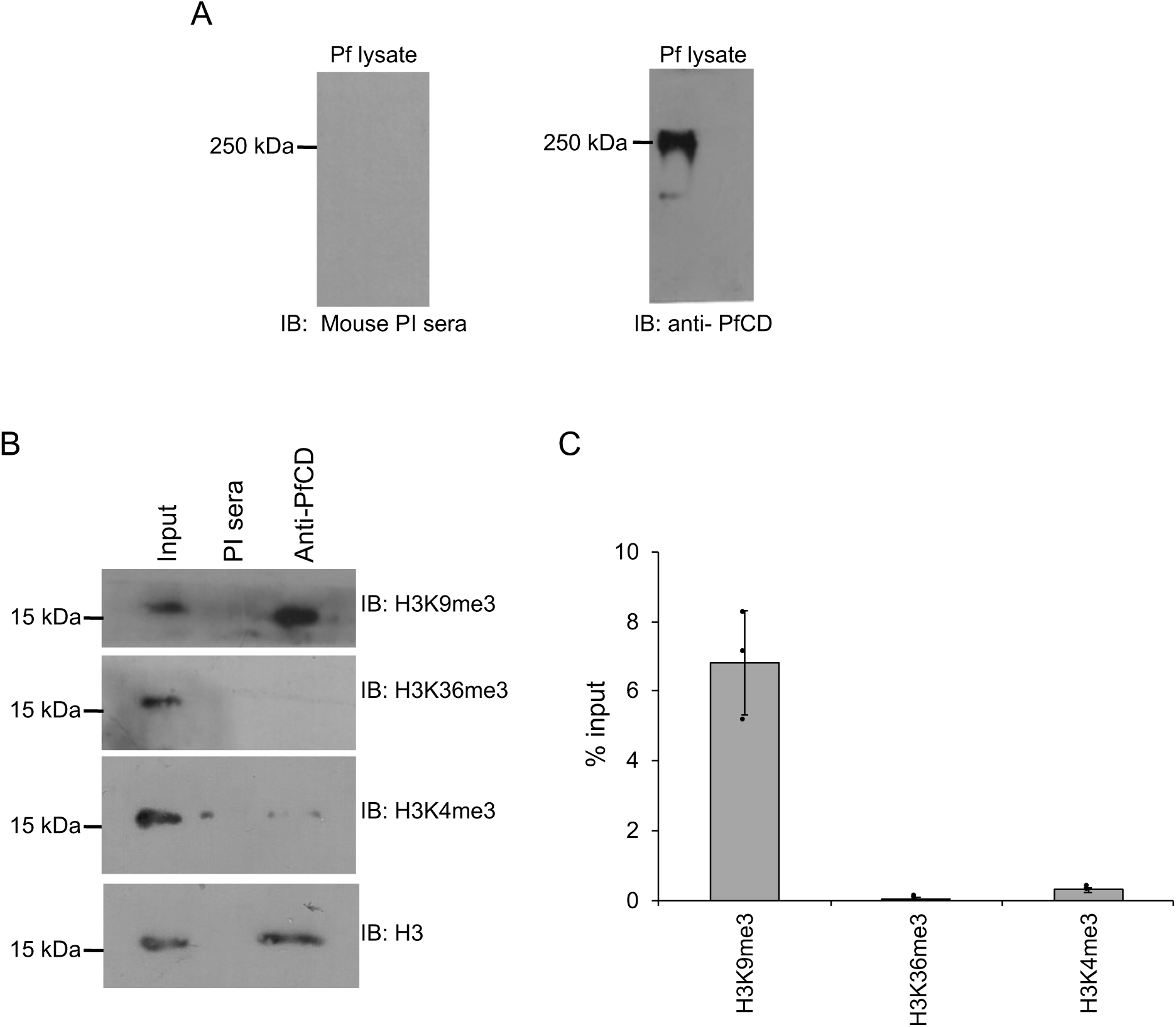
PfCDP interacts with H3K9me3 containing chromatin in *P. falciparum*. **(A)** Validation of anti-PfCD IgG for its PfCDP specific interaction with *P. falciparum* lysate. The mouse pre-immune sera were used as the control, does not react to *P. falciparum* lysate and anti-PfCD IgG specifically interacts to the ∼250 kDa protein matching to PfCDP in *P. falciparum*. **(B)** The PfCDP protein cross-linked chromatin from *P. falciparum* was prepared and immunoprecipitated using anti-PfCD IgG, and pre-immune sera were used as the control. The immunoprecipitated complexes were separated on 16% SDS-PAGE gel and probed with H3K9me3, H3K36me3 and H3K4me3 specific antibodies. In parallel, a separate ChIP reaction was performed, and subjected to immunoblot against H3 protein as a control for IP reactions. All the modifications ChIP reactions are performed in parallel and ChIP reactions were carried out in triplicate. The representative western blot images are provided. **(C)** The bar plot represents the band intensity of H3K9me3, H3K36me3 and H3K4me3 marks enriched in PfCDP – chromatin immunoprecipitation fraction. The band intensity was measured using ImageJ software and the error bar represents the SEM of three independent ChIP assays.

### Conditional deletion of chromo domain in PfCDP of *P. falciparum*

To explore the functions of the PfCDP – H3K9me3 interaction in *P. falciparum*, we used a recently developed Rapamycin inducible DiCre expression system in *P. falciparum* NF54 lines to generate a conditional gene deletion (34). The Rapamycin-mediated activation of DiCre recombinase has proven highly effective in mediating site-specific recombination and leads to the conditional deletion of target genes in *P. falciparum* (34, 35). We designed a construct with two loxP sites within artificial introns (LoxPint), each flanking the chromodomain, which underwent deletion upon the addition of Rapamycin. (Figure 4A; Supplementary figure 4A–4D). The correct integration of the artificial chromodomain exon flanked by the two loxPints was confirmed by PCR. The Rapamycin mediated excision of the chromodomain was confirmed by PCR using specific primers flanking the deleted region (Figure 4B). The Rapamycin induced deletion of the PfCD domain alone leads to a complete reduction of PfCDP at the transcript level in *P. falciparum* (Figure 4C). The western blot analysis with an anti-PfCD antibody against the chromodomain in the parasite lysate revealed the loss of PfCDP protein expression in Rapamycin treated *P. falciparum* (Figure 4D – 4E). Surprisingly, conditional deletion of the DNA encoding for the chromodomain alone leads to a complete reduction of the full CDP transcript and protein. One possibility is that the deletion of the CD encoding genomic DNA, which lies at the 5’ end of the gene, may lead to defects in the transcription factor assembly and failure in the transcription of the full-length gene.

**Figure 4:**
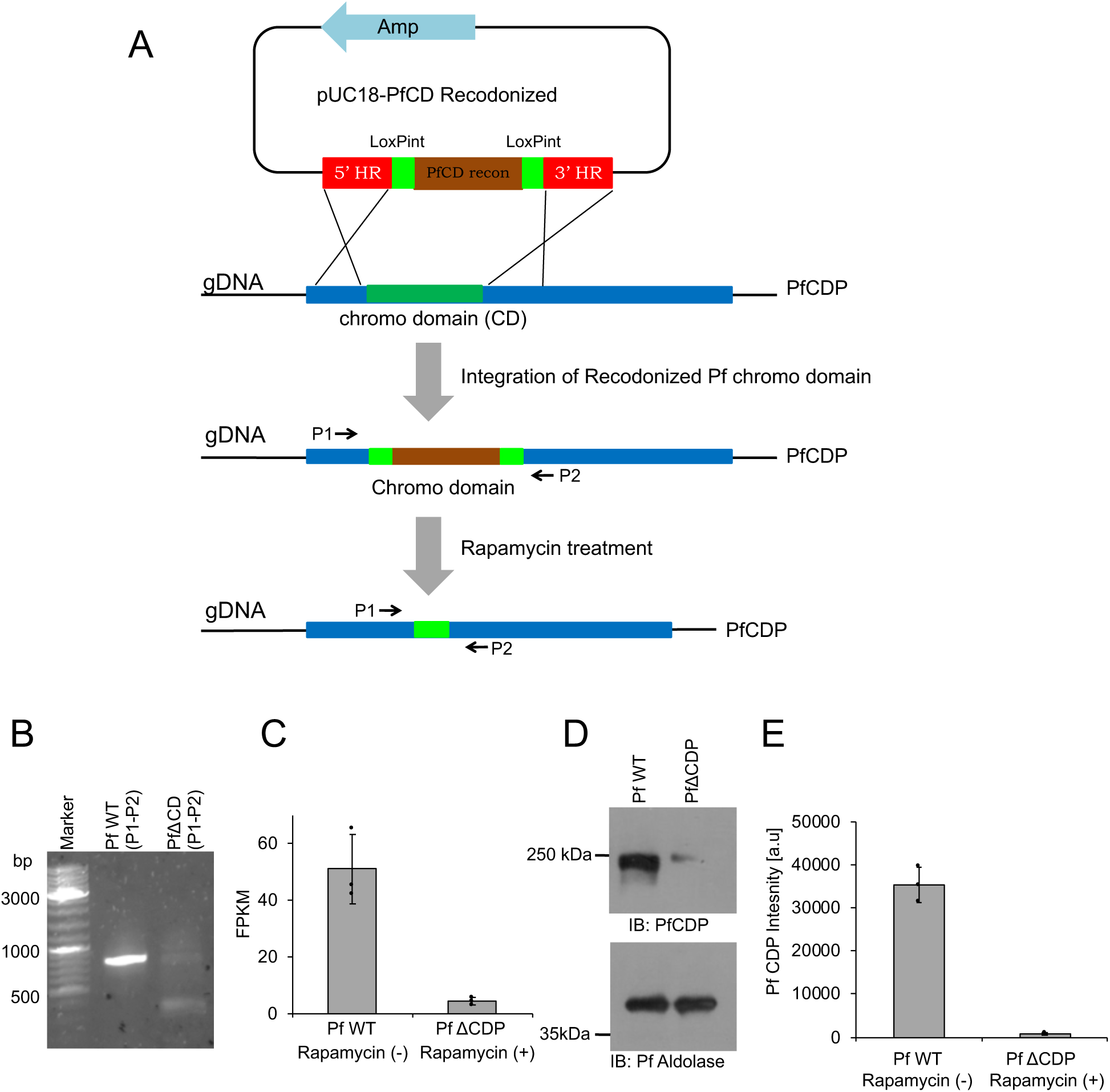
Conditional deletion of chromodomain (CD) in *P. falciparum*. **(A)** Schematic representation of the conditional deletion strategy for PfCD in *P. falciparum.* The pUC18 – PfCD vector was prepared using the recodonized chromodomain (Brown color) flanked with artificial loxPint (green color). The 5’ and 3’ regions flanking the chromodomain were cloned adjacent to loxPint (red color). The next scheme is a representation of CD (dark green color) in PfCDP (Blue color), and the cross line represents the homologous recombination through which the PfCD recodonized DNA will be integrated into the PfCDP genomic DNA. The P1 and P2 is the primer pair designed at homologous region (HR) sites. The bottom scheme represents the loss of CD fragment in the PfCDP gene after the Rapamycin treatment to *P. falciparum*. **(B)** PCR amplification with P1 and P2 primer pairs confirms the loss of CD in the genome of *P. falciparum* upon Rapamycin treatment than DMSO control. **(C)** The RNA sequencing data confirms the loss of PfCDP transcript in Rapamycin treated *P. falciparum,* and the error bar represents the SEM from three independent RNA sequencing data sets. **(D)** The immunoblot analysis using anti-PfCD antibody confirms the loss of PfCDP protein in PfΔCDP *P. falciparum* and bottom immunoblot for Pf Aldolase represents the loading control. **(E)** The bar plot represents the band intensity of PfCDP and the band intensities were measured using ImageJ software. The error bar represents the SEM of three independent biological immunoblots.

### PfCDP represses subset of *rifin* genes in *P. falciparum*

To understand the functional implication of PfCDP-H3K9me3 interaction on the regulation of gene expression in *P. falciparum*, we performed strand-specific RNA sequencing of the wild type and the mutant (PfΔCDP) lines of the parasite. The transcriptomics study was performed in triplicate for the *P. falciparum* wild type and PfΔCDP lines and obtained in high-quality sequencing reads (Supplementary figure 5) (Supplementary table 1) (**Supplementary file 1**). The differential gene expression (DGE) analysis validated the depletion of PfCDP expression in the PfΔCDP *P. falciparum* lines (DGE p-value =1.44E-73) (Figure 5A**; Inset Fig**). We have found that 19 genes were found to be upregulated commonly in all three PfΔCDP *P. falciparum* lines, of which more than 50% belongs to the virulence gene family members and the rest of the upregulated genes were mostly unspecified products including an apicoplast ribosomal protein L14 and a small subunit ribosomal RNA (Supplementary figure 6). We also observed 3 genes (including PfCDP) were found to be downregulated in the PfΔCDP *P. falciparum* line (Figure 5B).The Gene Ontology (GO) analysis of the differentially expressed genes revealed an enrichment of GO terms associated with biological processes like regulation of cell adhesion, antigenic variation, and cytoadherence to the microvasculature (Figure 5C). The *rifin* genes were abundant among the upregulated genes, in addition to a few *var* and *stevor* genes (Figure 5D). We did not observe any type-specific (Type A vs Type B RIFIN) enrichment among the upregulated *rifin* genes. To substantiate the up-regulation of subsets of *rifins* are specific effect due to PfCDP gene deletion, we have compared the expression levels of the *rifin* genes in wild type and under the PfCDP deleted condition. A significant number of *rifins* genes were expressed at higher levels upon PfCDP conditional deletion and these changes are specific effect due to the loss of PfCDP as we observed consistent up-regulation of subsets of *rifins* in CDP conditionally deleted *P. falciparum* lines (Supplementary figure 7). The functional implications of these up-regulated *rifins* in the development and progression of infection have not been addressed so far. However, clinical studies have reported the presence of antibodies against two of the upregulated *rifins* (Pf3D7_0300200 and Pf3D7_0833200) in *P. falciparum* severe malaria cases from malaria-endemic regions (43, 44). Thus, PfCDP mediated regulation of a subset of *rifin* genes may have a potential association with the development of *P. falciparum* severe malaria.

**Figure 5:**
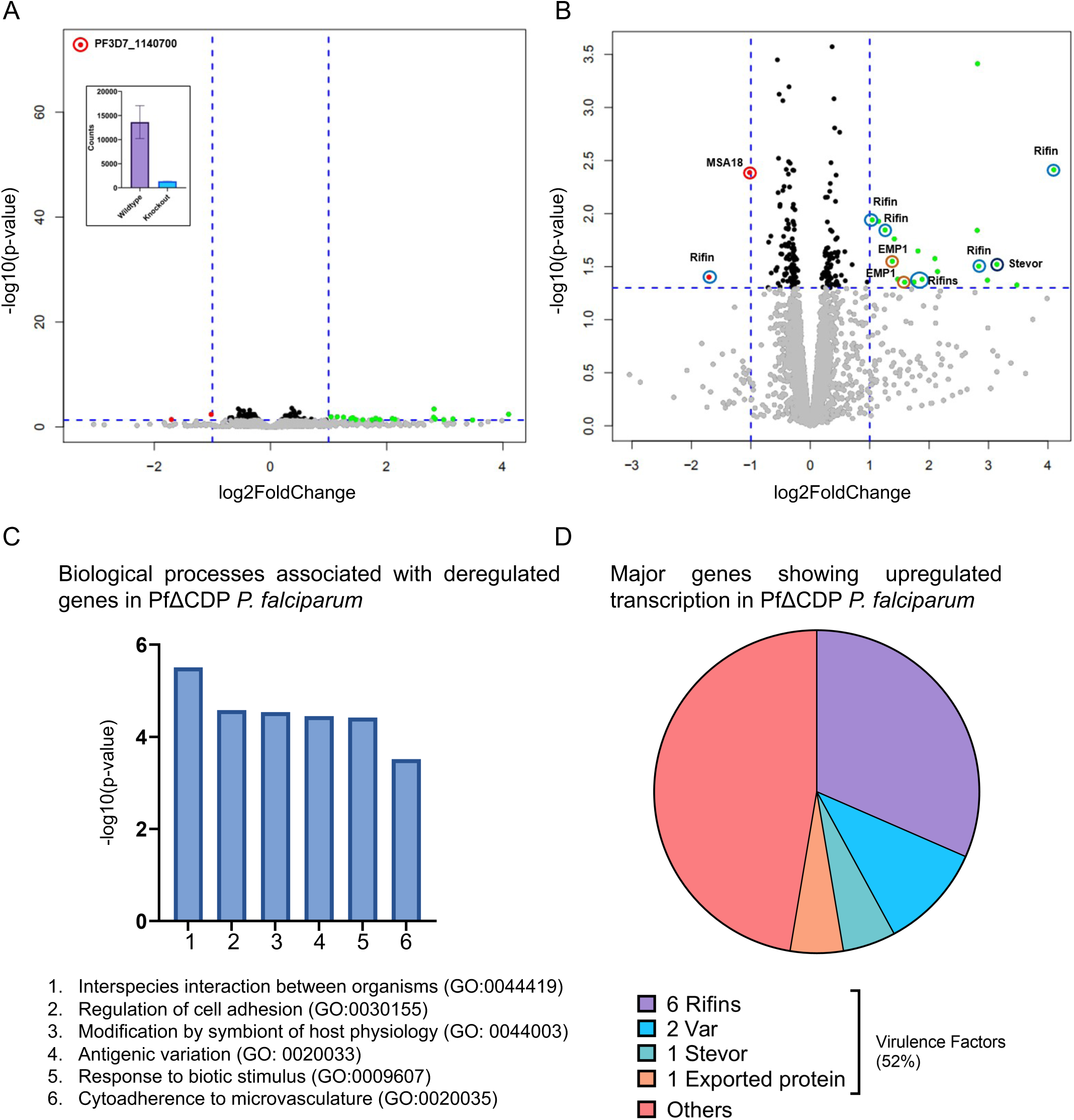
Investigation of transcriptional deregulation associated with the conditionally deleted PfCDP *P. falciparum* lines. **(A)** Volcano plot representing the differentially expressed genes in PfΔCDP vs wild type *P. falciparum*. The PfCDP gene (PF3D7_1140700) shows the most statistically significant deregulation among all genes (p-value at 1.44E-73). **Inset image:** A histogram representing the depleted read counts of PfCDP in the conditional deleted parasite lines compared to wild type. **(B)** Volcano plot representing the statistically significant (p-value <=0.05) 19 up-regulated (green) and 3 down-regulated (red) genes in PfΔCDP vs wild-type *P. falciparum*. A significant number of deregulated genes are virulence-associated factors (*Rifin*, *Var*/EMP1, and *Stevor*). Note: The PfCDP gene (down-regulated with p-value =1.44E-73) has been removed from the visualization to prevent skewing of the plot. **(C)** GO Terms and IDs of biological processes were found to be deregulated significantly in the PfΔCDP *P. falciparum* lines as inferred by Gene Ontology (GO) analysis. **(D)** The pie chart represents the genes that are significantly deregulated genes in PfΔCDP *P. falciparum* calculated from 3 independent biological replicates of RNA sequencing.

### The PfCDP binds to subsets of *rifins* genes and regulates its expression in *P. falciparum*

To study the mechanism of PfCDP mediated regulation of *rifins* genes, we performed the ChIP sequencing for PfCDP protein to map its positioning on the global genome scale in *P. falciparum*. The PfCDP ChIP sequencing gave us 857 peaks of enrichment mapped on 693 genes with >= 1.8 fold enrichment over and above the input (Figure 6A) (**Supplementary file 2**). For a vast majority of the target the occupancy of PfCDP was found at the entire gene body starting from transcription start site (TSS) to the transcription end site (TES) (Figure 6A). Further, we observed that the PfCDP1 occupancy was depleted from both the gene flanking 5’ and 3’ UTR regions. Around 35% of the target genes of PfCDP were found to be associated with cytoadherence and antigenic variation associated functions. These include 121 *rifins*, 57 *var* and 24 *stevor* genes (Figure 6B) (**Supplementary file 3**). The Gene Ontology enrichment analysis of the target genes gave high enrichment of cytoadherence associated biological functions (Figure 6C). The epigenetic heterochromatin mark H3K9me3 was also found to be highly enriched on these loci (Figure 7A**) (**Figure 7C**) (Supplementary** figure 8) (Supplementary figure 9). Notably, both PfCDP and H3K9me3 modification occupancy were found to be depleted on non-virulence gene loci (Figure 7B). Next, we performed a correlation analysis of the global PfCDP1 and H3K9me3 genomic occupancy and discovered a moderate correlation between the two with a spearman correlation of coefficient, R=0.43 (Figure 7D). This correlation is driven strongly by the absence of both the elements from non-virulence loci and moderately by their enrichment from virulence gene clusters (Figure 7D).

**Figure 6:**
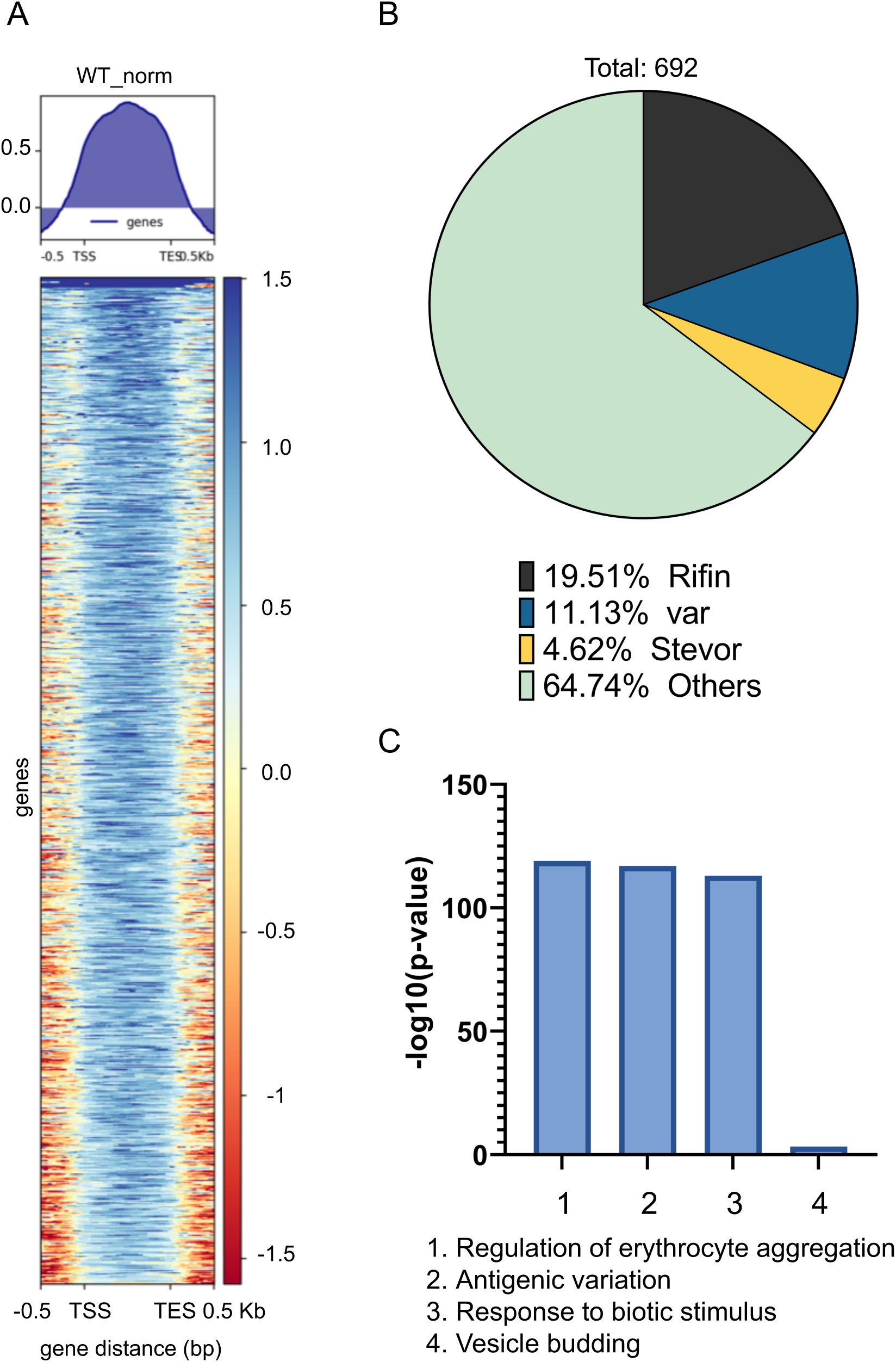
PfCDP is enriched upon gene targets predominantly associated with virulence functions in *P. falciparum*: **(A)** Heatmaps representing the enrichment of PfCDP on the gene bodies of target genes with >=1.8fold enrichment. **(B)** Pie charts representing the proportion of the 692 genes that are virulence associated genes (*var, rifin, stevor*) vs others. The *rifins* are the major targets among the virulence genes to the PfCDP in *P. falciparum* **(C)** Bar plots representing the strong enrichment of PfCDP targets for biological functions associated with virulence, cytoadherence and antigenic variation.

**Figure 7:**
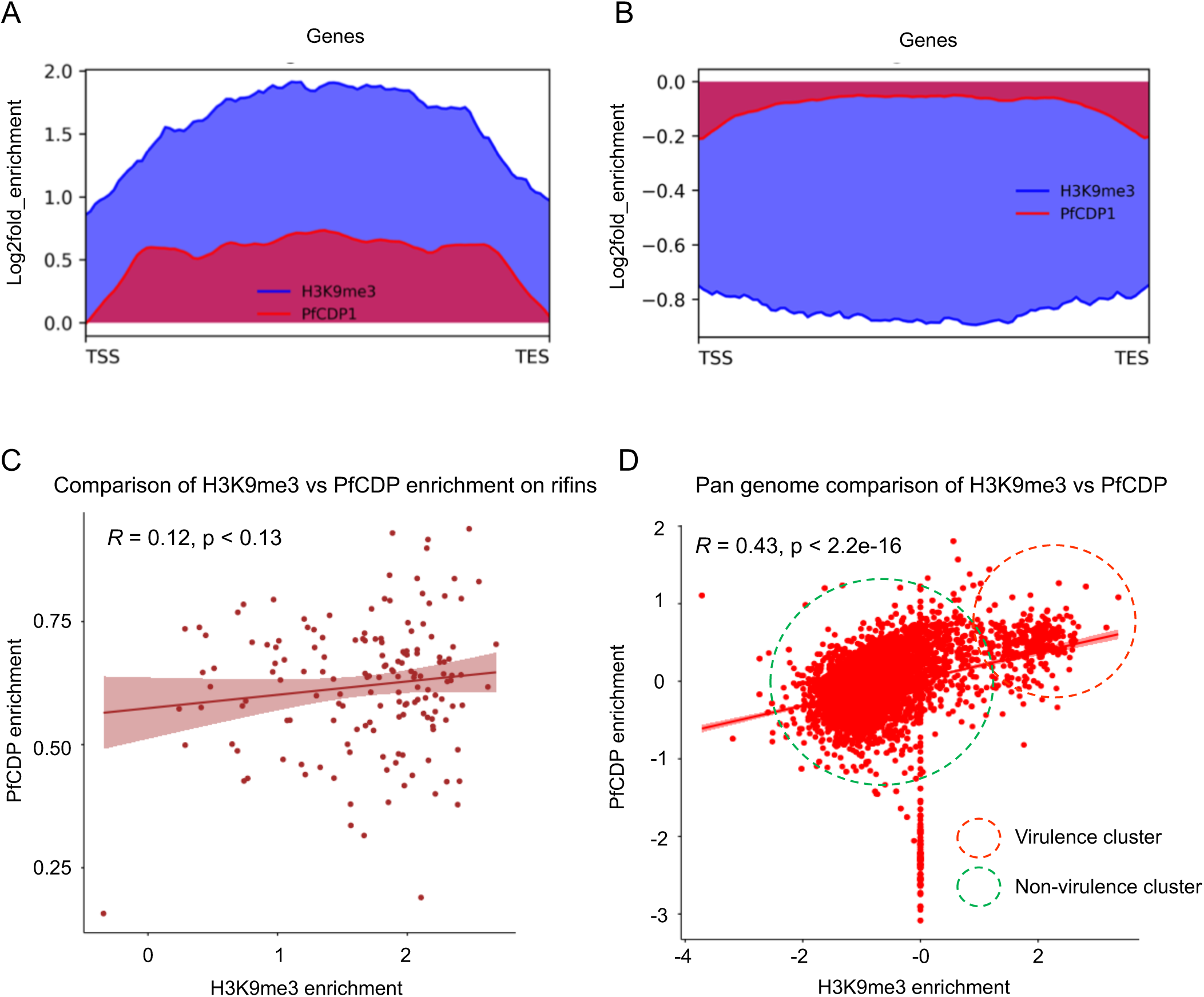
PfCDP is enriched upon heterochromatic, virulence associated gene clusters: **(A)** Strong enrichment profile of PfCDP (red) and the heterochromatin marking histone modification H3K9me3 (blue) on the average gene body in virulence associated gene clusters. **(B)** Both PfCDP and H3K9me3 are depleted over the non-virulence associated genome in the parasite. **(C)** Scatter plot comparing the co-existence of PfCDP and H3K9me3 modification on the rifin gene subset. **(D)** Scatter plot representing the pan genomic distribution of PfCDP and H3K9me3 with a positive correlation driven by strong depletion of both from non-virulence genes and moderate enrichment of PfCDP on virulence genes.

We checked for the correlation of PfCDP occupancy with the expression of the its target genes and discovered that compared to all genes, the PfCDP target genes have a lower average expression (Figures 8A – 8B). This is an indicative of a gene repressive environment associated with PfCDP occupancy. This is further supported by the fact that PfCDP correlates with heterochromatin mark H3K9me3, a strong gene silencing modification in *P. falciparum*. The virulence genes like *var, rifins and stevors* are members of multigene families which show mutually exclusive expression, which requires a majority of them to be kept silenced. PfCDP1’s strong occupancy on these loci and its association with H3K9me3 modification may contribute to their highly controlled expression.

**Figure 8:**
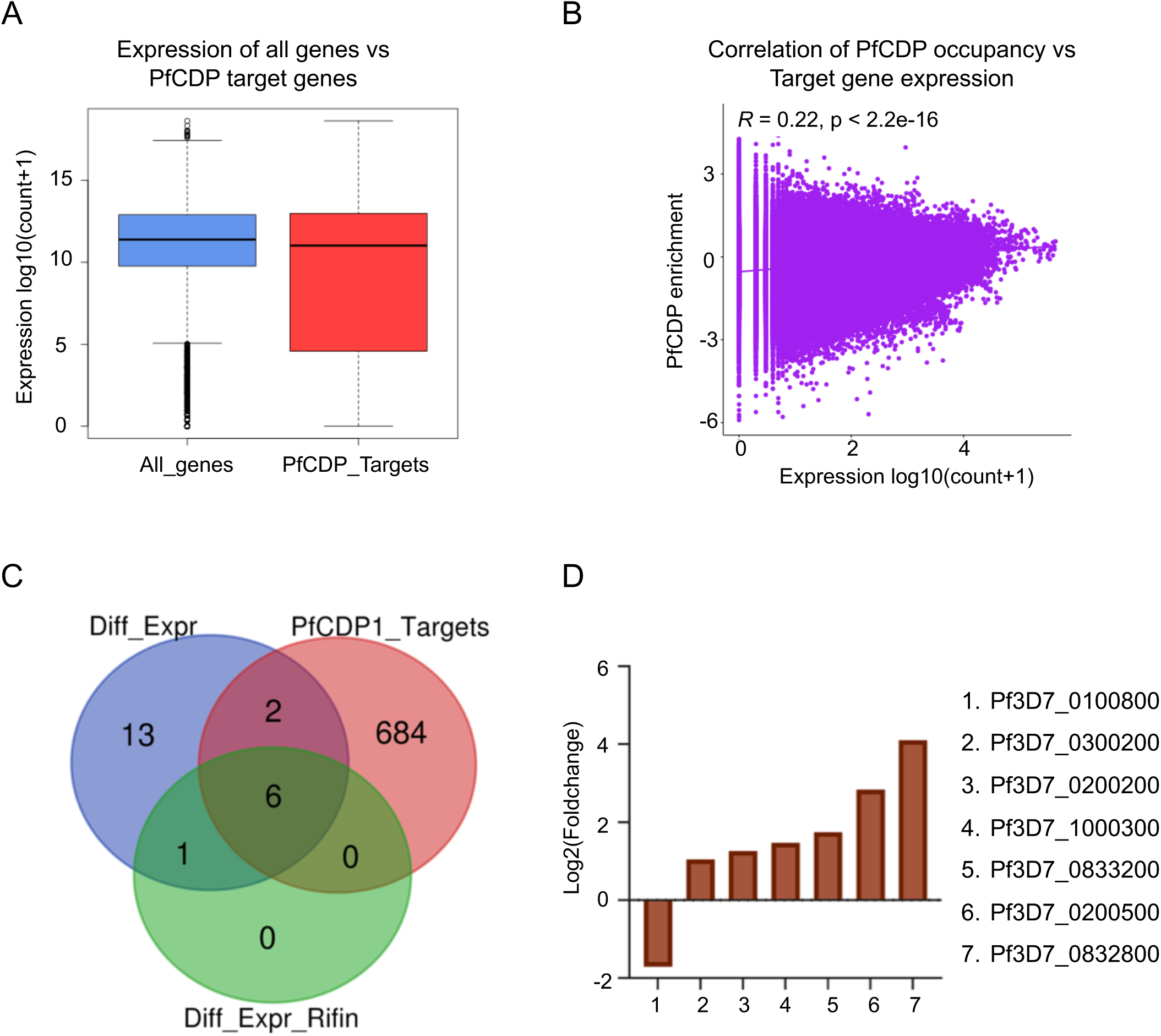
PfCDP occupancy is neutrally correlated with the expression of a large fraction of the parasite genome: **(A)** Boxplots comparing the average expression profile of all genes vs PfCDP enriched genes shows an overall similar mean expression. The expression of a number of PfCDP target genes (virulence genes) is however lower than the average. **(B)** Scatter plot comparing the normalized read counts for PfCDP occupancy vs gene expression over pan genomic bins shows little to no correlation between the two. **(C)** Venn diagrams comparing the overlap of PfCDP targets (red) with differentially expressed genes in PfCDP conditionally deleted *P. falciparum* line (blue) and differentially expressed rifins (green). More *rifins* with barely threshold enrichment of PfCDP seem to deregulate upon its deletion. **(D)** The expression values from RNA sequencing experiments in conditionally deleted CDP lines have confirmed that the observed top 6 upregulated *rifins* are the major PfCDP targets in *P. falciparum*.

We evaluated the expression profile of all genes as well as the virulence associated genes *var, rifin and stevors* under PfCDP wildtype and PfCDP deletion condition and discovered littles to no deregulation of total transcriptome and *var* gene sets (Supplementary figures 10A – 10B). Importantly, we observed that the *rifins* have a moderately higher expression upon PfCDP deletion (Supplementary figure 11A), and we did also observe a higher average expression of all *stevor* genes under PfCDP deletion (Supplementary figure 11B). Next, we have analysed the number of differentially expressed genes are the target of PfCDP1 and found that 8 out of 22 differentially expressed genes (Supplementary figure 6) are the direct targets of PfCDP (Figure 8C). The transcriptome analysis with conditionally deleted PfCDP strain shows deregulation of 22 genes including a subset of virulence genes (3 downregulated and 19 upregulated); out of these 6 *rifins* were found to be upregulated upon PfCDP deletion in *P. falciparum* (Figure 5D). We analysed whether these upregulated *rifins* are the targets of PfCDP and found all the 6 *rifins* shows enrichment of PfCDP (Figures 8D). Although, the ChIP sequencing for PfCDP yielded 693 target genes, it has high positive correlation only to the differentially expressed virulence genes, particularly to the subsets of *rifins* in *P. falciparum* and these targets are concomitant with the heterochromatin repressor epigenetic mark H3K9me3. In conclusion, the conditional deletion of PfCDP leads to up-regulation of subsets of *rifins*, which suggest that PfCDP involves in the regulation of only small percentage of *rifins* genes in *P. falciparum*.

### Conditional deletion of the chromodomain in PfCDP promotes RBC rosette formation by *P. falciparum*

The functions of the majority of the RIFINs are not studied; however, recent reports suggest its involvement in RBC adhesion and rosetting (31, 45). Rosetting is an important virulence mechanism that contributes to the development of severe malaria in humans by *P. falciparum* (46). Rosettes are formed by the recruiting uninfected RBCs towards the *P. falciparum*-infected RBCs, adapted by the parasite to escape the immune system. Further rosettes block the capillaries that eventually lead to the development of severe malaria (22, 31). In addition to RIFINs, other virulent proteins such as EMP1 and STEVOR contribute to rosette formation (19, 47). We observed the up-regulation of a subset of RIFINs in PfΔCDP *P. falciparum* lines and therefore tested whether these RIFINs can induce rosette formation *in vitro*. The *P. falciparum* wild type and PfΔCDP lines were cultured in O^+^ RBCs and confirmed the loss of the PfCD domain by PCR in Rapamycin treated parasites (Figure 9A) and we collected the rosette RBCs from the culture. We observed 7-10% rosettes in *P. falciparum* wild type lines; whereas, significant levels of increased rosettes with PfΔCDP lines than wild type *P. falciparum* (Figure 9B, Supplementary figure 12). The estimation of rosette formation has revealed that conditional deletion of PfCD in the parasite promotes rosettes by three-fold compared to the wild type *P. falciparum* lines (Figure 9C). The observation of increased rosettes in PfΔCDP *P. falciparum* lines establishes a strong correlation with the up-regulated subset of RIFINs and suggests the potential involvement of the virulent proteins in the process of RBCs rosetting.

**Figure 9:**
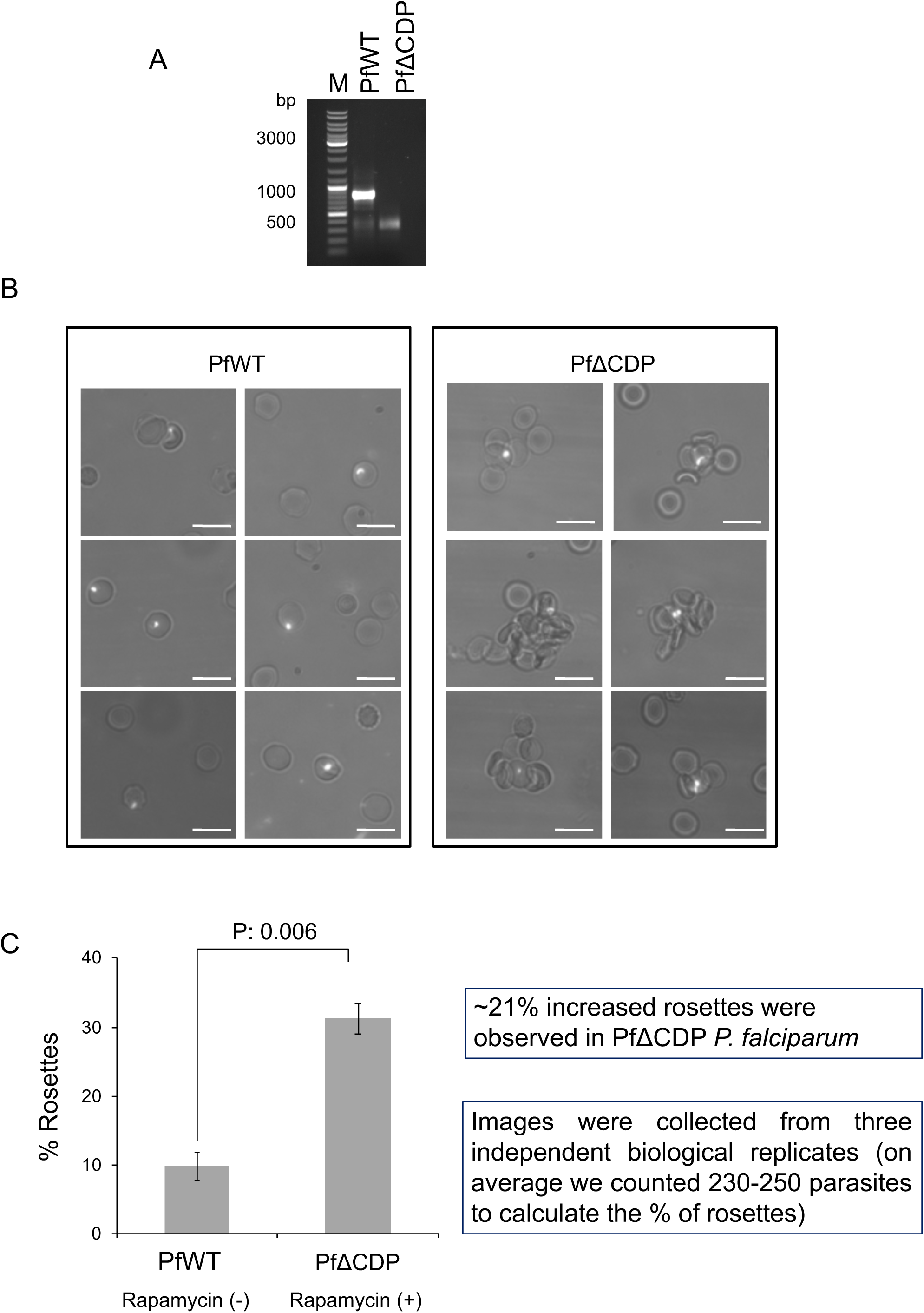
PfΔCDP *P. falciparum* promotes the RBC rosetting. **(A)** Agarose gel image of PCR amplicons confirms the loss of CD gene fragment in the Rapamycin treated *P. falciparum* culture batch used for rosette assay. In every biological replicate assay, the loss of CD fragment was verified and collected the rosettes from the *P. falciparum* culture. **(B)** Representative microscopic images of *P. falciparum* rosettes, increased rosettes were observed in PfΔCDP *P. falciparum* than wild type lines. The rosettes were collected from *P. falciparum* cultured in O^+ve^ RBCs and the parasite stained with Hoechst 33342 nuclear stain. The images were obtained using an Olympus BX61 microscope fitted with a 60X objective lens. The *P. falciparum*-infected RBCs are shown in the dense white spot due to the parasite nuclear material stained with Hoechst 33342. The scale bar represents 10 μm. The additional RBC rosette images are provided in supplementary figure 5. **(C)** The bar plot represents the percentage calculation of rosettes from wild type and PfΔCDP *P. falciparum* lines. The percentage of rosettes were calculated from 230 – 250 parasites counted from each biological replicate experiment using the formula: Number of rosettes / Total number of mature parasites × 100. The error bar represents the SEM from three independent rosettes assays. The calculated p values were significant and calculated using the paired t-test.

## Discussion

The severity of malaria differs among individuals and primarily depends on antigenic variation mechanisms employed by *P. falciparum* (22, 48, 49). Major proteins that exhibit antigenic variation include PfEMP1, RIFINs, and STEVORs. They are expressed by the parasite, and many are exported to the surface of infected RBCs (22, 50, 51). The proteins are encoded by multigene families by approximately 60 *var*, 150-200 *rifins*, and 30-40 *stevor* genes (6, 19, 32, 52, 53). The expression and regulation of the *var* genes are well studied and tightly regulated by two epigenetic marks H3K9me3 and H3K36me3 in *P. falciparum* (23–25, 47, 54). The comprehensive global mapping of epigenetic marks has revealed major virulence genes such as *var, rifins, stevors, and Pf-TM* are enriched with the H3K9me3 mark (Lopez-Rubio et al., 2009). The HP1 chromodomain is the first epigenetic reader protein characterized in *P. falciparum* that interacts with the H3K9me3 mark and regulates the expression of *var* genes, parasite invasion, phenotypic variation, and gametocytogenesis (24, 26, 38). Here we identified a second chromodomain-containing protein that interacts with the H3K9me3 mark in *P. falciparum* and controls the expression of a subset of RIFINs and rosetting of RBCs but only a few *var* genes and no genes associated with gametocyte development (Figure 5D; Figure 10). The selective up-regulation of only a subset of RIFINs indicates that additional mechanisms are likely important for RIFINs regulation, or, that many are expressed in other lifecycle stages.

**Figure 10:**
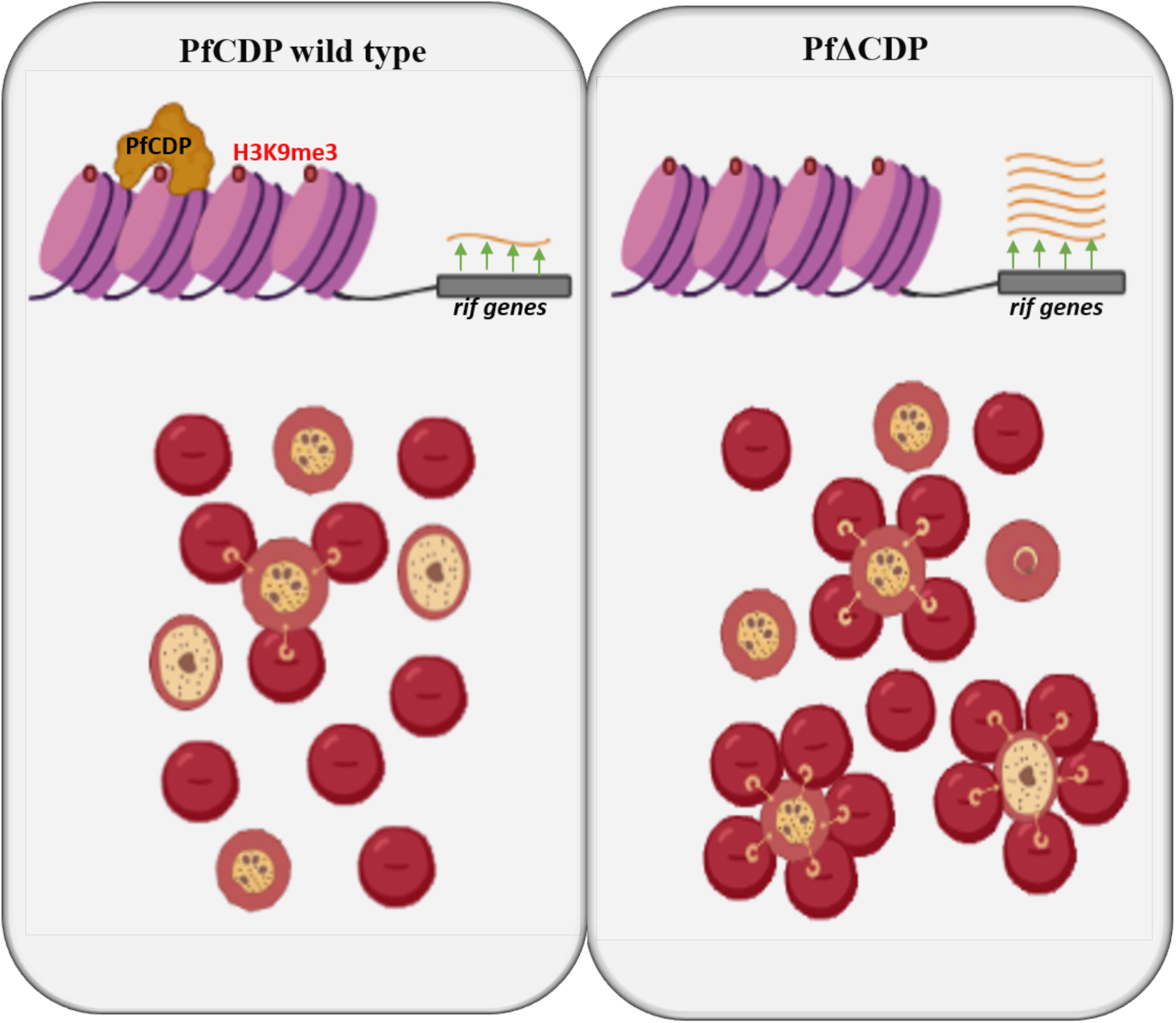
Schematic representation of PfCDP mediated regulation of RIFINs expression and rosetting of RBCs through H3K9me3 modification specific interaction on the chromatin of *P. falciparum.* The deletion of PfCDP promotes the RBCs rosetting, which implies the significance of the RIFINs in the virulence mechanisms in *P. falciparum*.

The ChIP-sequencing for PfCDP had indicated that it binds to multiple *rifin* targets, however, only subsets of *rifins* are deregulated upon conditional deletion of CD in *P. falciparum.* The *P. falciparum* genome contains approximately 200 copies of *rifins*, and the PfCDP does not bind to all the *rifins,* which indicates that the additional chromodomain containing proteins or other chromatin binding protein might be involved in controlling the expression of the complete set of *rifins*. The PfCDP majorly binds to the *rifins* targets and some extend to the *stevor* and *var* genes and regulation of these multigenes in *P. falciparum* seems a very complex process. A key question is: how does PfCDP control a subset of *rifins* selectively? Or more general: what drives the specificity of epigenetic readers that control the *var* gene expression, *rifins*, or genes required for sexual stage differentiation? The recent identification of the gametocyte development protein 1 (GDV1) gene to specifically control sexual stage development by removing HP1 from heterochromatic loci (55) indicates a complex level of protein-protein interactions that control the specificity of methyl reader proteins. It is also possible that antisense RNA or long-non-coding RNA regulatory mechanisms are involved in controlling the virulence mechanisms through altering the epigenetic network in *P. falciparum* (25, 56).

Rosetting of RBCs in malaria patients is a hallmark for severe malaria, and recent reports have identified that the rosetting is mediated through virulence proteins, particularly RIFINs and PfEMP1 (31, 46) The rosetting mediated development of severe malaria is highly variable among individuals and one of the host factors that dictate the outcome of the malaria severity of ABO blood group antigens (31, 57). Goel *et al* have reported that one of the RIFINs (PF3D7_0100400) mediates the rosetting of RBCs, primarily with A blood group RBCs (31). Another study has shown that two RIFINs (PF3D7_1254800 and PF3D7_0223100) interact with the leucocyte-associated immunoglobulin-like receptor 1 (LAILR1) on B- and NK cells to inhibit their activation, thereby suppressing the development of immune response against parasites in the host (30). A clinical study has shown the presence of antibodies to four RIFINs among infected individuals (43, 58).

The subset of RIFINs up-regulated upon deletion of PfCPD leads to a significant increase of rosetting, indicating that one or several of them may play an essential role in complicated malaria. Antibodies to the RIFINs (Pf3D7_0300200 and Pf3D7_0833200) have been detected in severe malaria patients in malaria-endemic areas (43) (44). Both the genes exhibited significant up-regulation upon the deletion of PfCDP in our study, suggest that the PfCDP regulates clinically relevant antigens in *P. falciparum*. Whether individual RIFINs are responsible for the observed increase in rosettes in our study or if it is a combination of them is currently not known and will need to be investigated in the future.

## Data availability and materials

All the data generated or analysed during this study are included in this article and its supplementary information files. The NGS files related to RNA sequencing are generated in this study are available for download from NCBI GEO accession number: GSE158585. The ChIP sequencing NGS files are available for download from NCBI GEO accession number: GSE180668.

## Ethics statement

Human RBCs used in this study were collected from healthy volunteers’ blood samples, with due approval from the institutional (RGCB) ethics committee (IHEC/01/2014/05). Healthy volunteers provided written consent to collect blood at RGCB Molecular Diagnostic facility.

## Acknowledgement

This work was supported by DBT – IYBA (Grant # BT/08/IYBA/2014) to AR. The authors acknowledge the University Grants Commission, Government of India for the senior research fellowship of VSD (UGC/Dec 2014/ 371086), CAJ (UGC/Dec 2015/365030) and GG is supported by Senior Research Fellowship, from Indian Council for Medical Research. VSD acknowledges the EMBO-short term fellowship program (Fellowship Number 7799). KK acknowledges support from the Genome Engineering Technologies program (BT/PR25858/GET/119/169/2017) of the Department of Biotechnology, Government of India. The authors acknowledge Dr. Moritz Treeck, The Francis Crick Institute, UK, for the help on generation of conditional CD knock-out in *P. falciparum*. We acknowledge Dr. Rakesh Laishram, Scientist EII, RGCB for sharing his lab facilities during this critical pandemic period to complete the experiments successfully. We also like to acknowledge Dr. Jackson James, Scientist F, RGCB, for helping with the microscope facility. We extend our thanks our friends who carefully edited the manuscript.

## Conflict of interest

The authors declare that they have no conflict of interest with the contents of this article.

## Author contributions

AR conceived the study, designed the experiments and supervised the project. VSD and AR performed the research, CAJ, GG performed nucleosome extraction, in vitro GST pull-down assays. MT helped in preparing the KO constructs and optimized the transfection of the plasmids and integrating the recodonized CD into the genome of *P. falciparum*. AK and KK analysed the RNA sequencing and ChIP sequencing data, VSD and AR analysed the experimental data. AK and KK drafted the RNA seq and ChIP seq results and methods. AR and VSD jointly prepared the manuscript draft. All the authors read and approved the final manuscript.

---

**Supplementary Figure 1:**
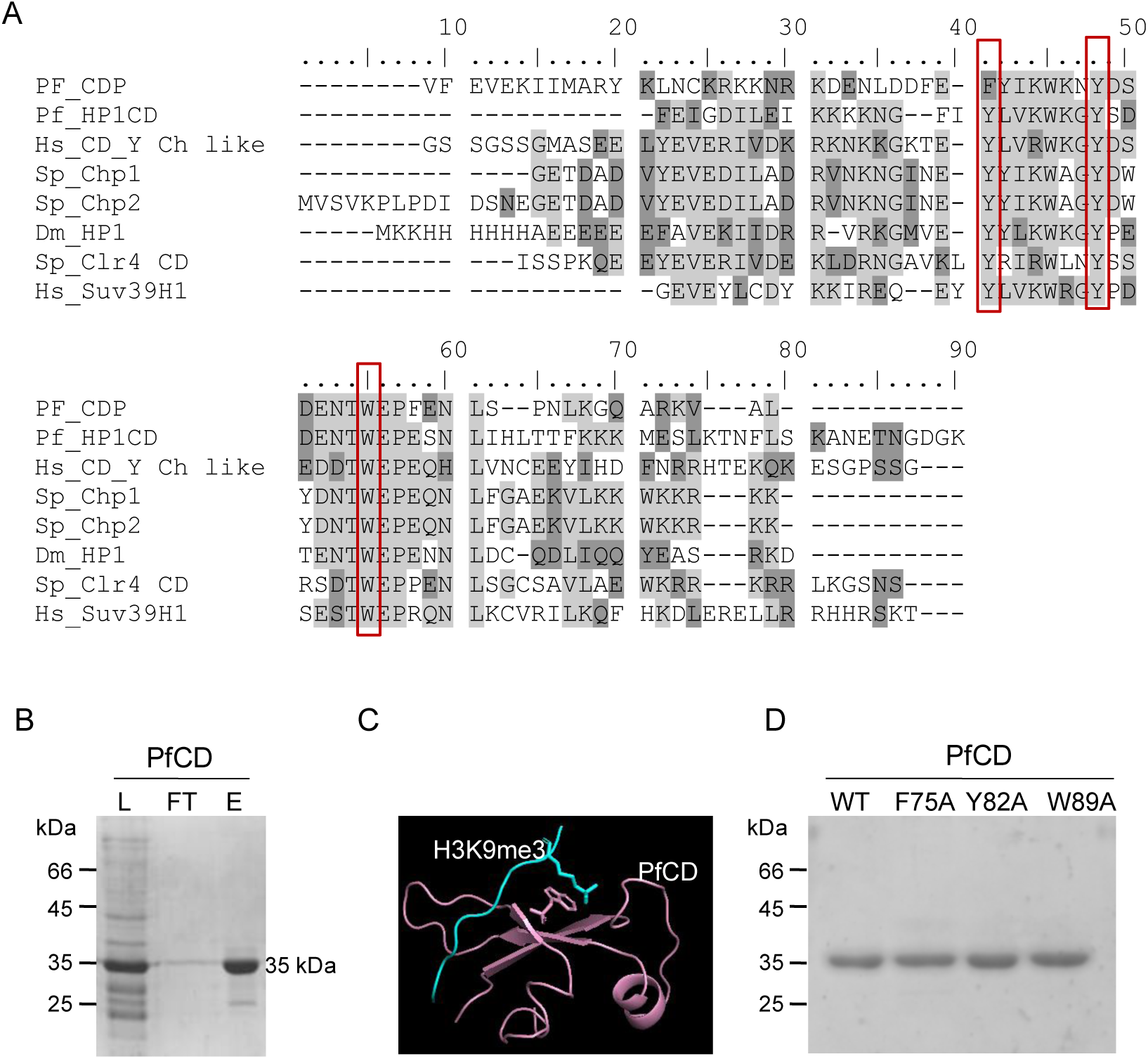
(A) Sequence homology analysis of Pf chromodomain with well-characterized chromodomain from other eukaryotic organisms that interacting with the H3K9me3 mark. *Pf: P. falciparum, Hs: H. sapiens, Sp: S. pombe, Dm: D. melanogaster.* The red colour box indicates the conserved aromatic amino acids that form the methyl binding pocket were selected for the mutational study. **(B)** Expression and purification of the PfCD domain as GST fused recombinant protein. L: Load, FT: Flow-through, E: Eluted fractions. Protein samples were separated on 12% SDS-PAGE gel and stained with Coomassie brilliant blue. **(C)** Homology model generation for PfCD and docking with H3K9me3 peptide to identify the potential aromatic amino acids pocket. H3K9me3 peptide is represented in blue colour. **(D)** Purification of PfCD Wild type and mutant proteins (F75A, Y82A, and W89A) as GST fused recombinant protein. Protein samples were separated on 12% SDS-PAGE gel and stained with Coomassie brilliant blue.

**Supplementary Figure 2:**
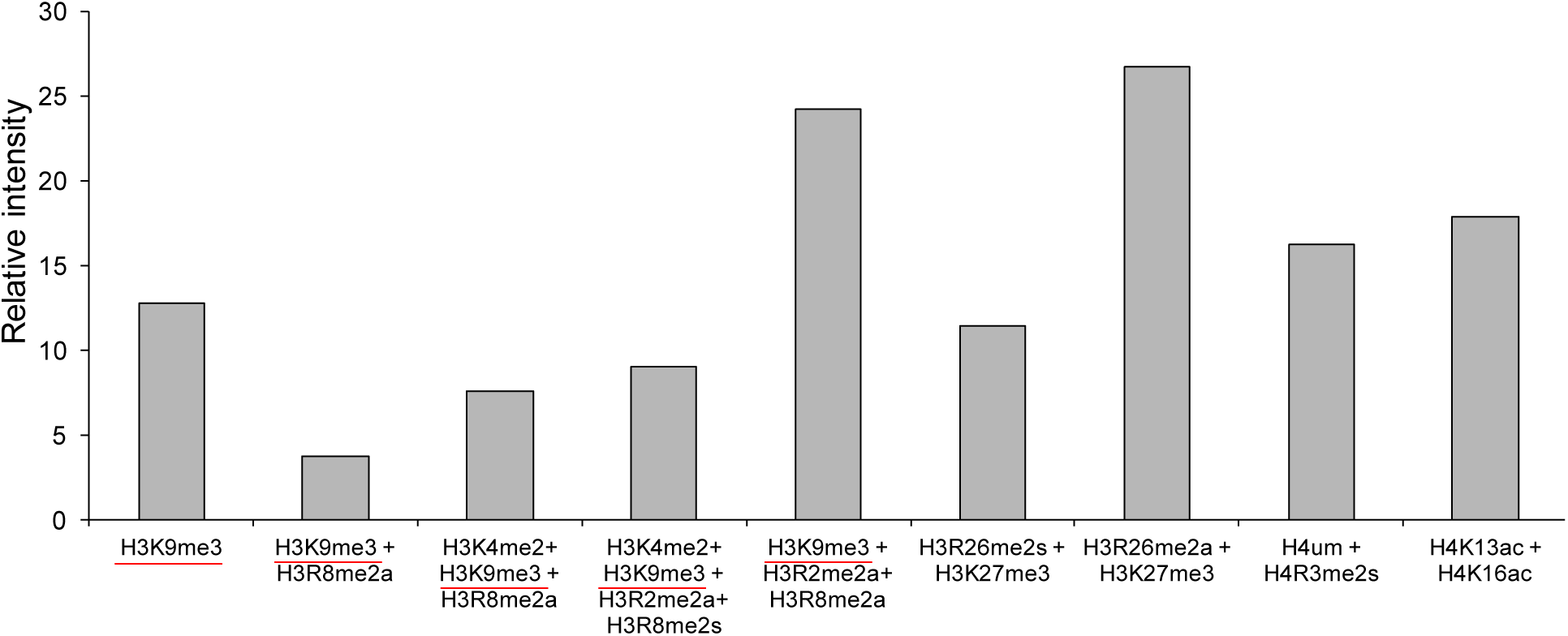
The peptide array spot intensity analysis was carried out using ImageJ and the relative intensity was calculated by normalizing to the intensity signal from unmodified peptides. The red colour underlines represents the H3K9me3 modifications in theses peptides along with other co-modifications.

**Supplementary Figure 3:**
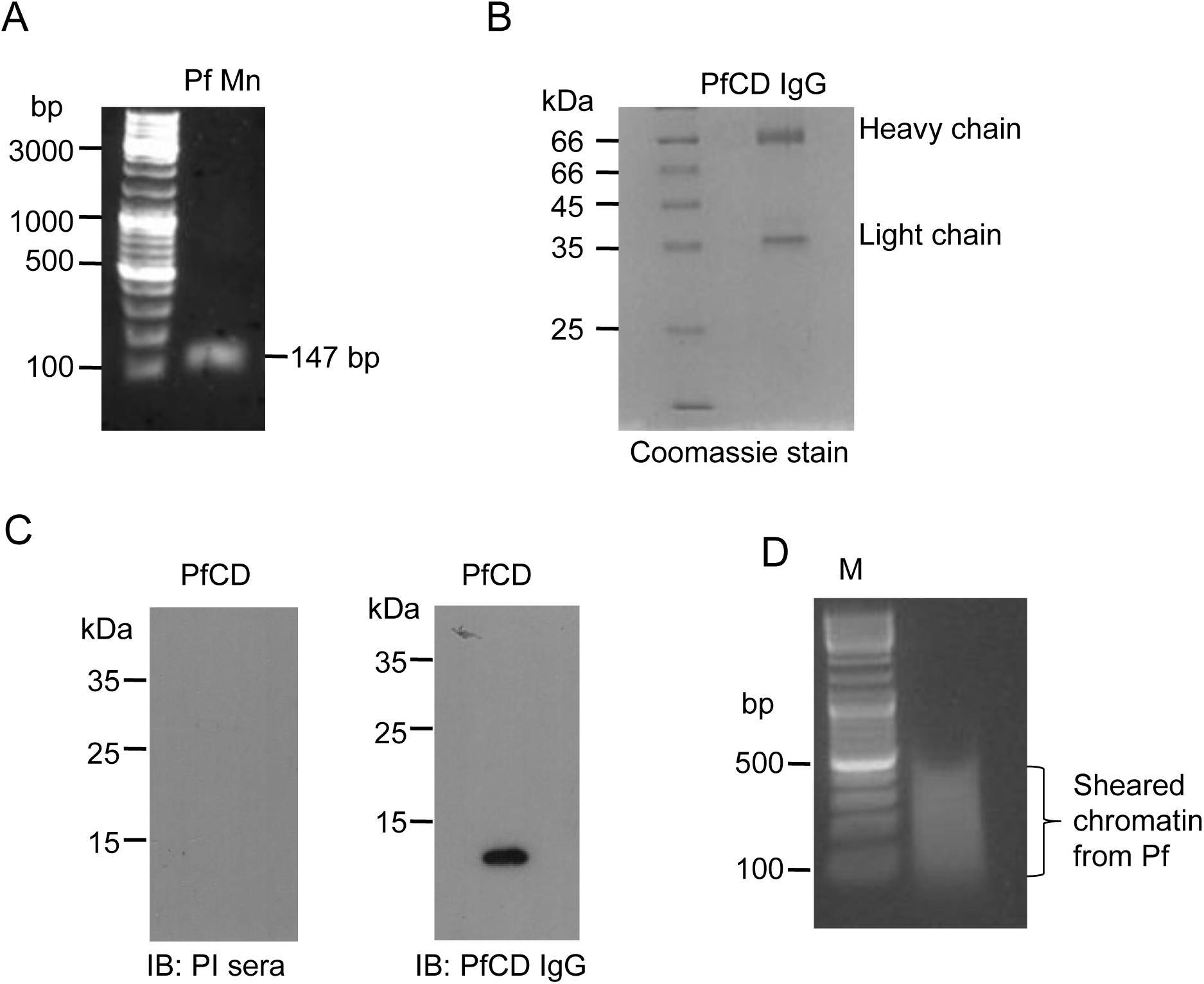
**(A)** Representative agarose gel image of mononucleosomes prepared from *P. falciparum* trophozoite stages (Pf Mn: *P. falciparum* mononucleosomes). **(B)** The purified PfCD IgG fraction from immunized serum collected from mouse and separated on 12% SDS-PAGE gel stained with Coomassie BB, the heavy and light chains of immunoglobulin are marked. **(C)** The western blot analysis of GST cleaved recombinant PfCD domain with purified PfCD IgG confirms the antibody specificity, and the pre-immune (PI sera) sera does not react to the recombinant protein. **(D)** Agarose gel image represents the quality of the separated sheared chromatin prepared from formaldehyde fixed *P. falciparum* for ChIP assays.

**Supplementary Figure 4:**
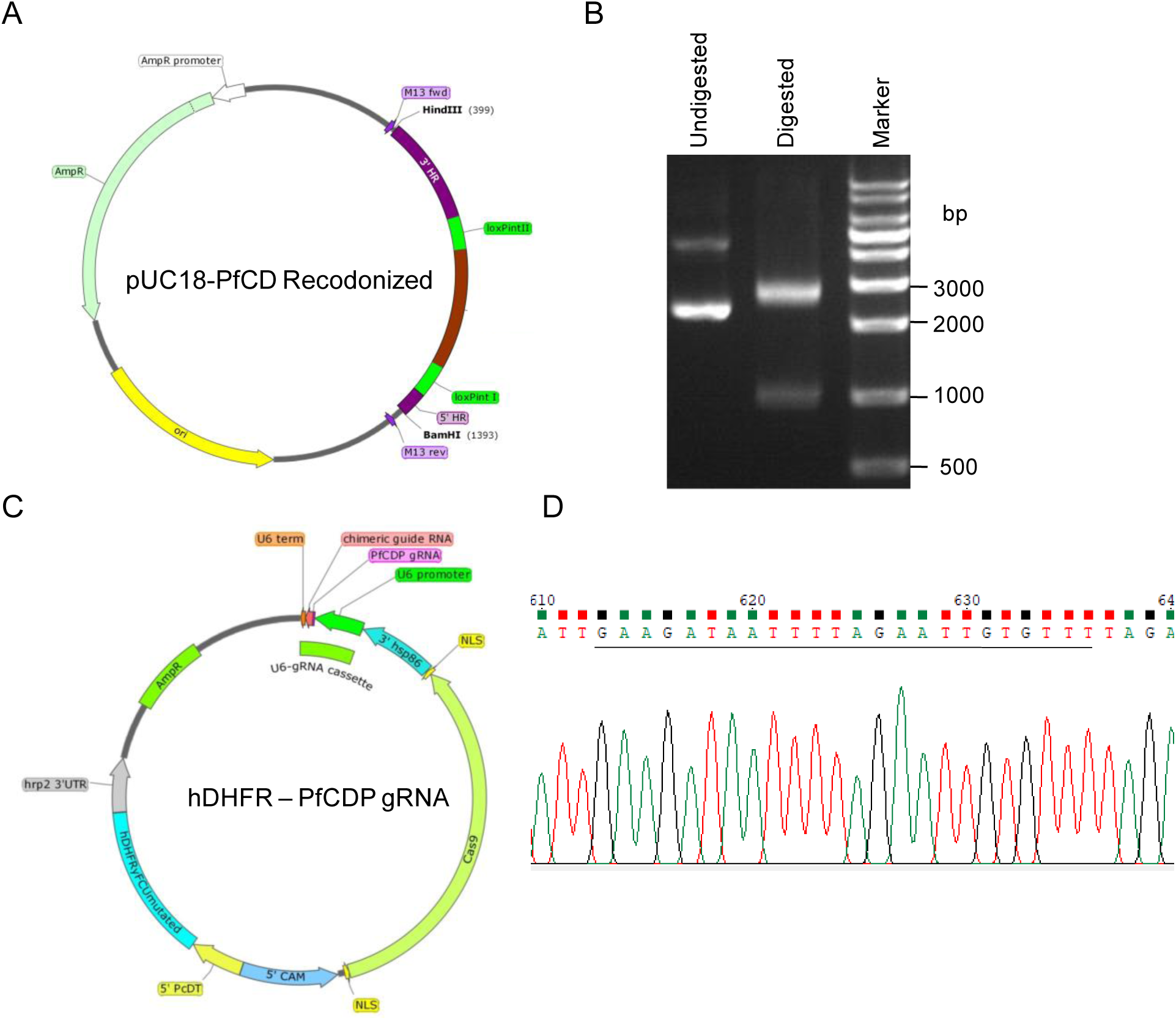
**(A)** Schematic representation of the PfCD recodonized vector map. **(B)** Restriction digestion pUC18-PfCD recodonized vector with HindIII, and BamHI enzymes confirm the release of the expected amplicon size product. **(C)** Schematic representation of the PfCDP gRNA vector map. **(D)** Sanger DNA sequencing confirms the presence of gRNA in the vector; the underlined sequence represents the gRNA.

**Supplementary figure 5.**
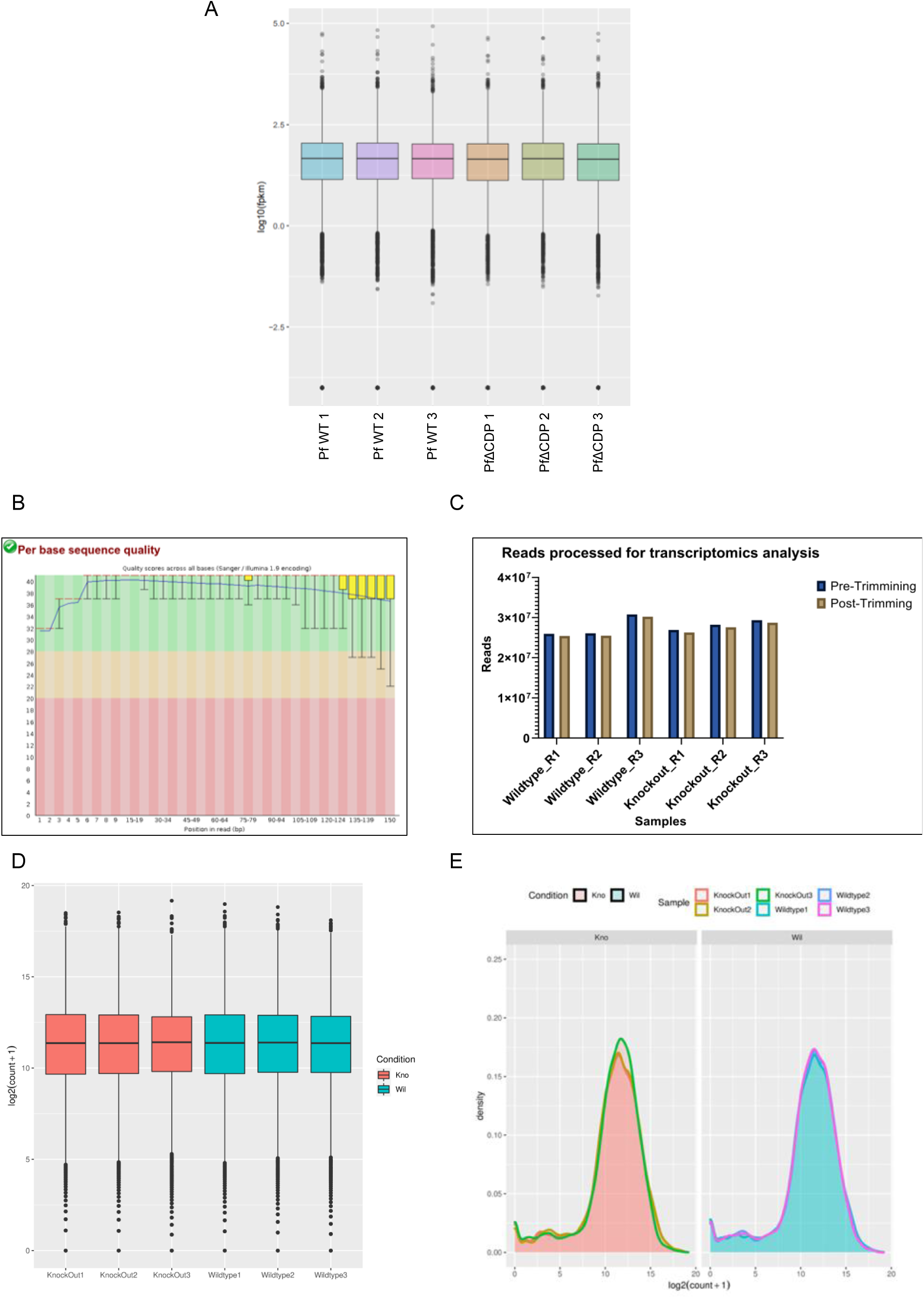
Analysis of transcriptional deregulation associated with knockout of PfCDP (Transcriptomics Quality Control Images). **(A)** The box plot represents the fragments per kilobase per million reads mapped (FPKM) for three wild type samples and three PfΔCDP samples. (**B**) A FastQC assessment file depicts the high quality of reads (phred score > 30) obtained with the RNA sequencing. **(C)** Histogram representing the number of reads processed (close to 25 million per sample) by Trim Galore! (high base-call scores and adaptor trimming) for further analysis. **(D)** A boxplot for the normalized read counts distribution across wildtype and knockout samples. **(E)** A density plot for the normalized read counts distributed across the wildtype and Knockout samples

**Supplementary figure 6.**
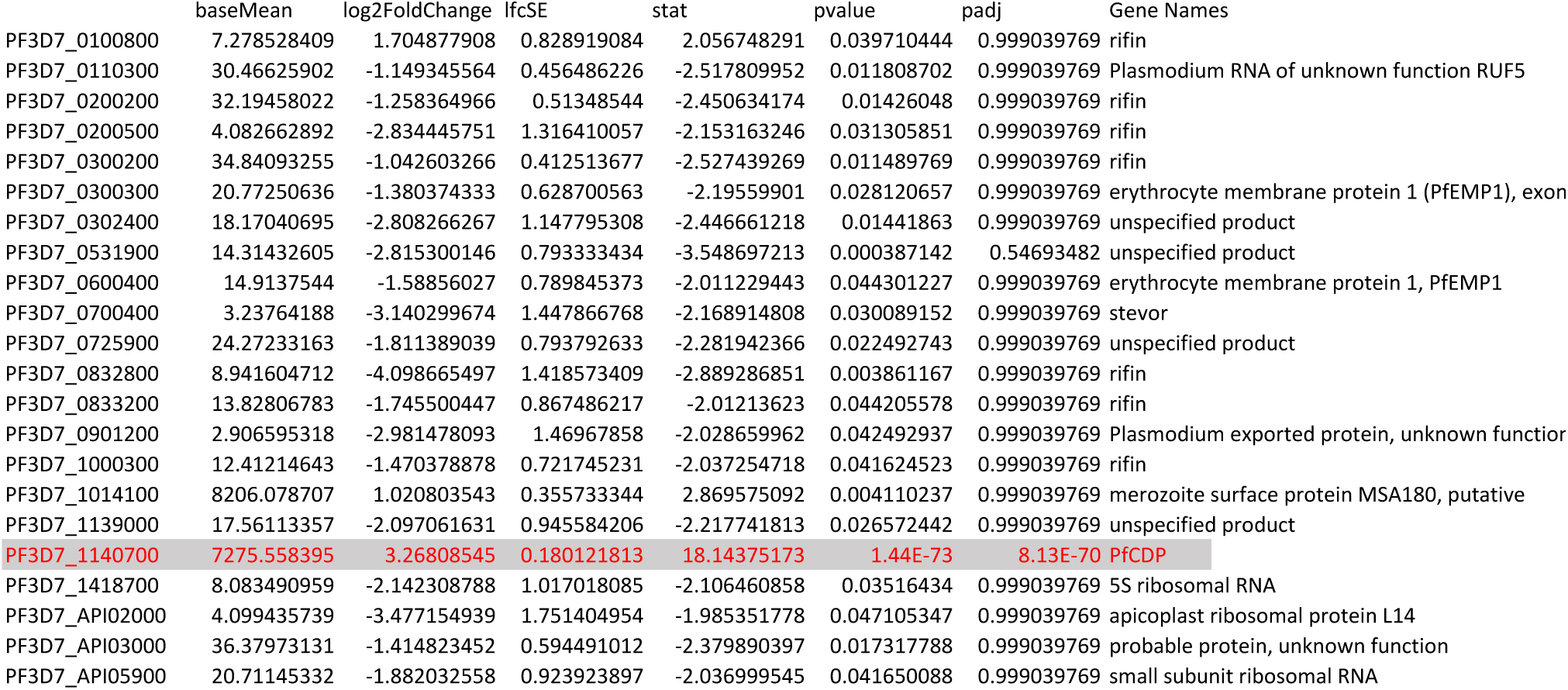
The list of genes found to be differentially expressed with Log2FC </>=1 and p value <0.05 in PfΔCDP *P. falciparum*. The deleted target gene (PfCDP) is highlighted in red color.

**Supplementary figure 7:**
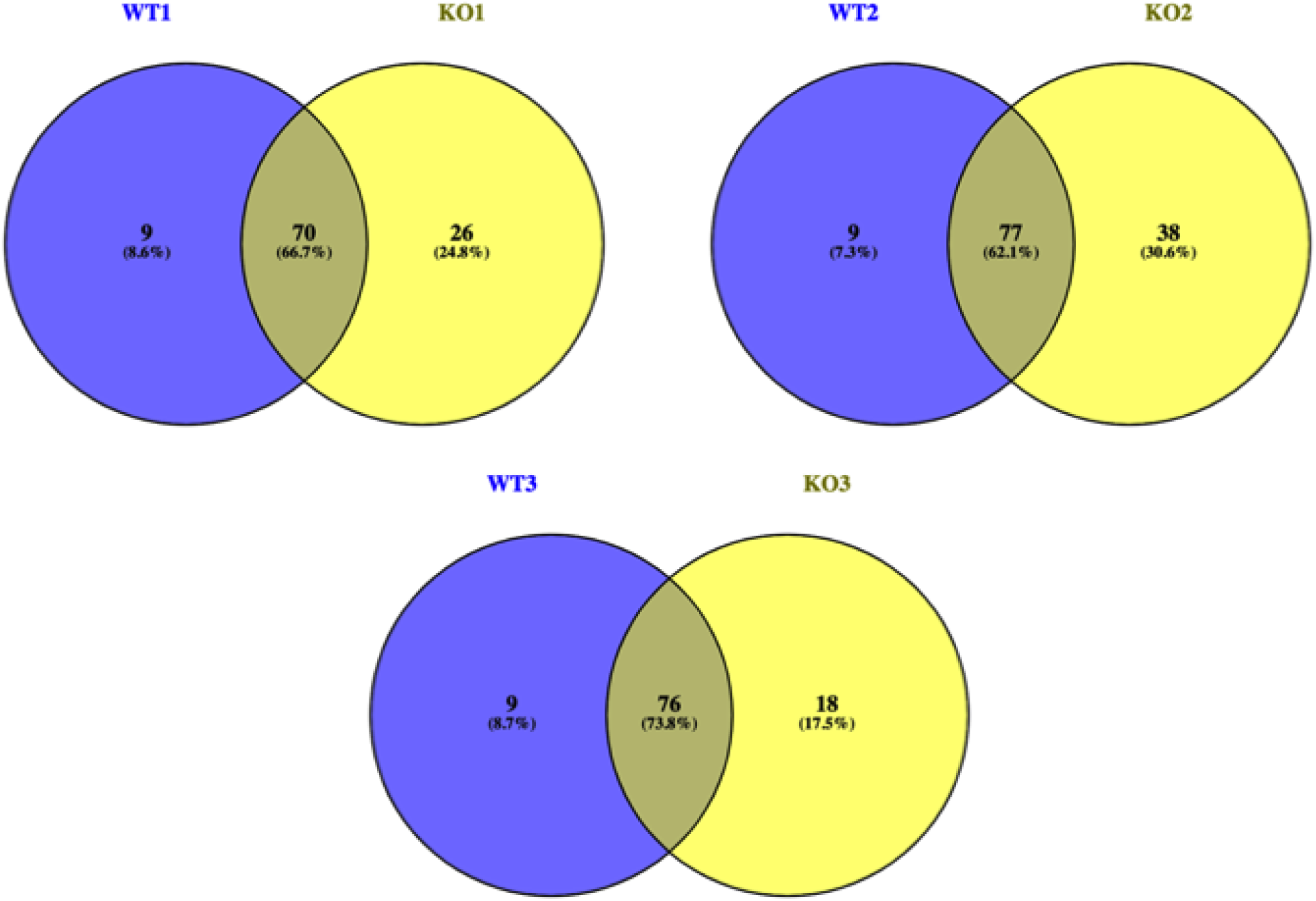
Venn diagram representing the *rifin* genes which are expressed (>10 read count) in WT1 vs KO1, WT2 vs KO2 and WT3 vs KO3 samples. Significant number of rifins are upregulated in CDP KO *P. falciparum* stain.

**Supplementary figure 8:**
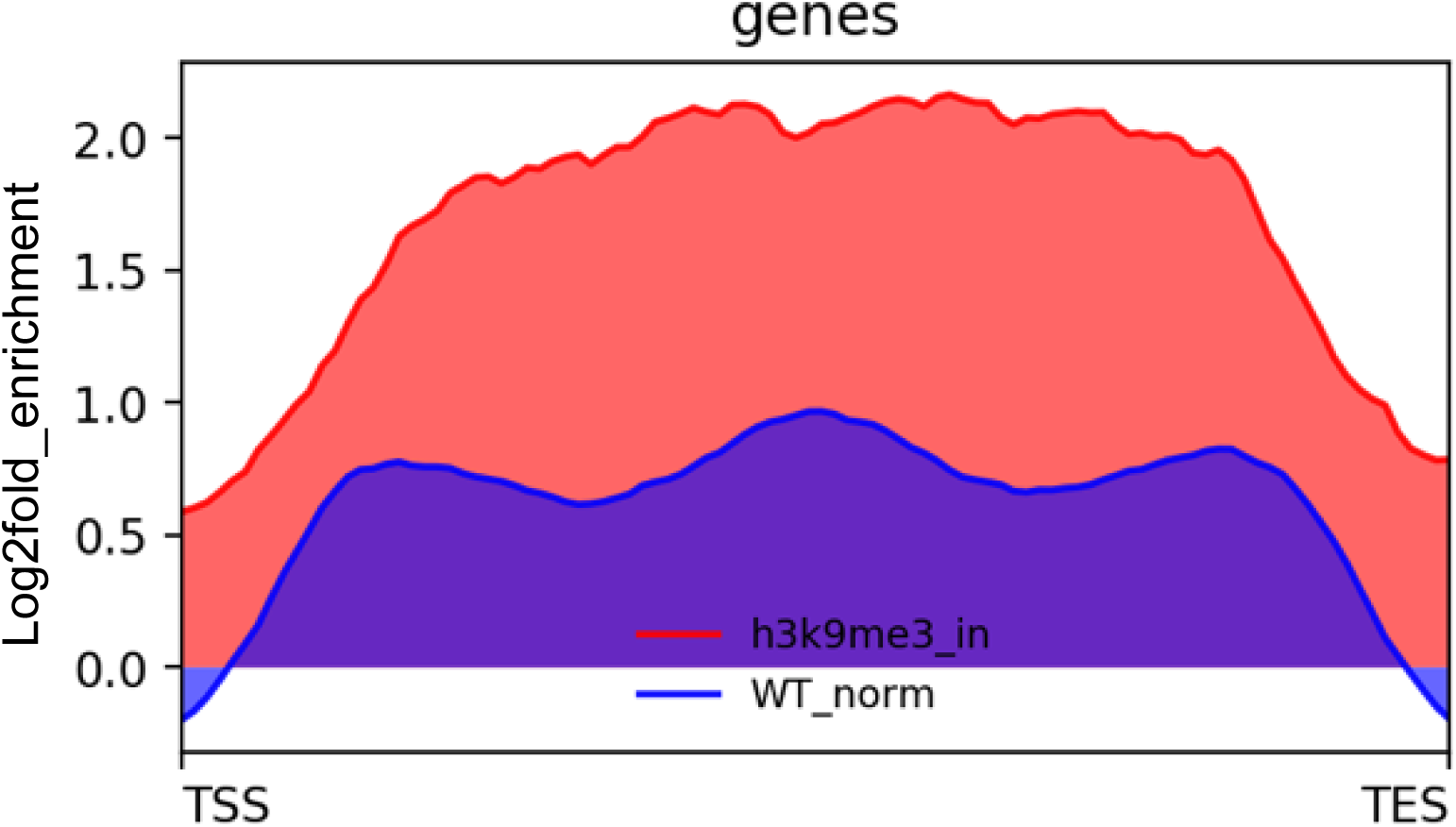
PfCDP enrichment on the H3K9me3 associated gene clusters, we detected good enrichment of PfCDP on the epigenetic heterochromatin mark H3K9me3 mark.

**Supplementary figure 9:**
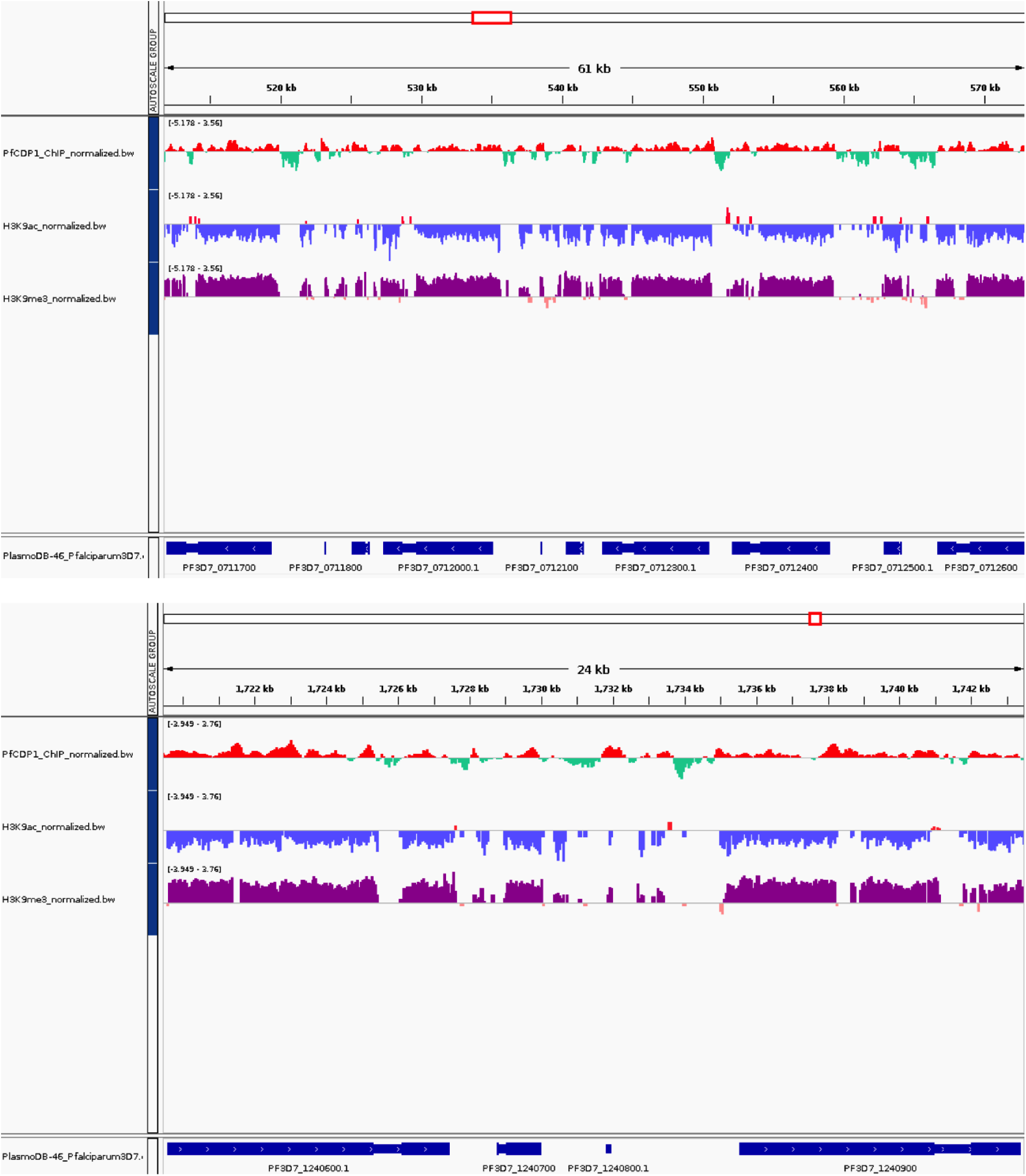
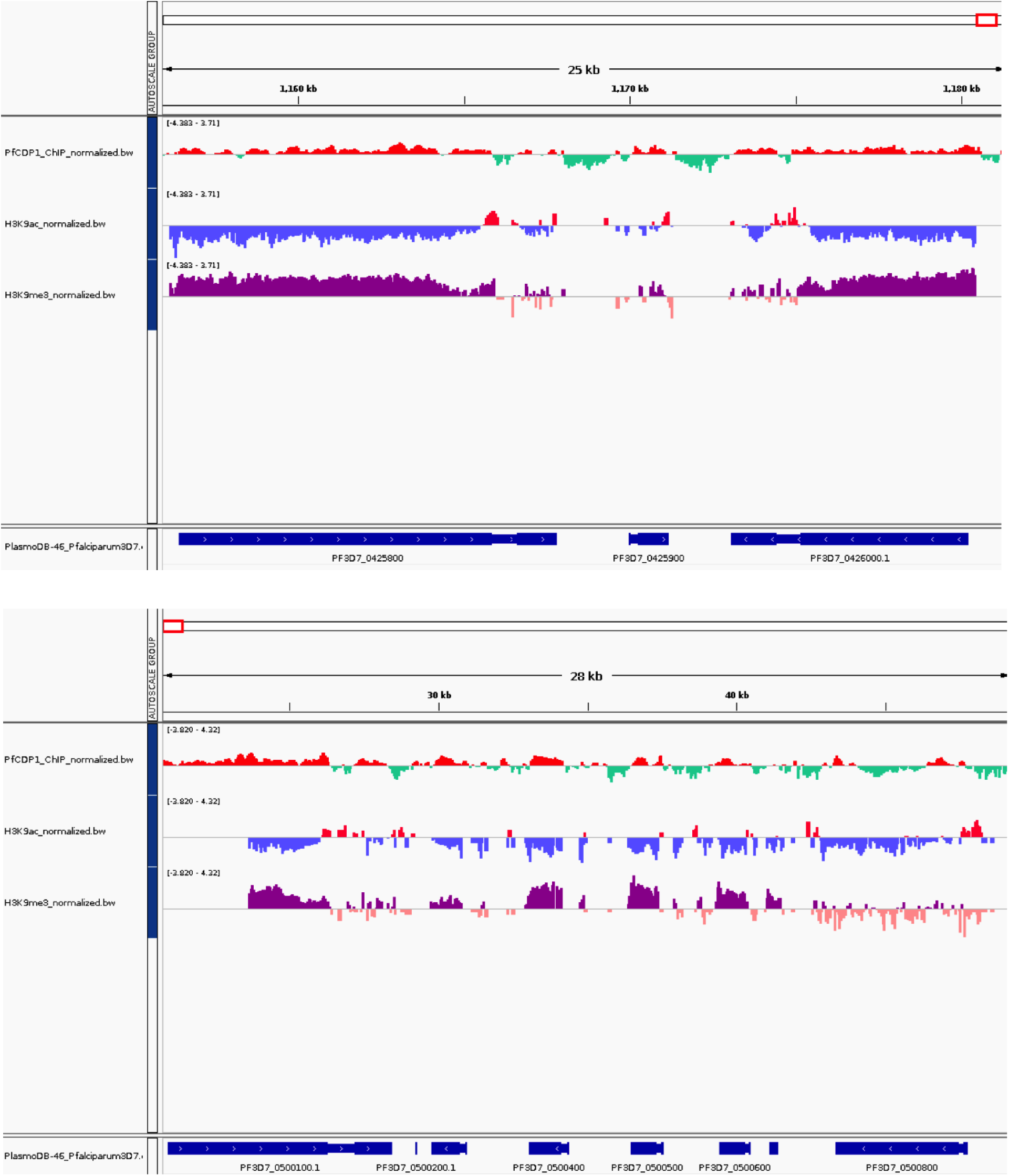
Representative chromosome track images for PfCDP ChIP sequencing reads with selected virulence genes. There is high a correlation of virulence genes repressor methylation mark H3K9me3 with PfCDP, whereas negative correlation of gene activation mark H3K9ac.

**Supplementary figure 10:**
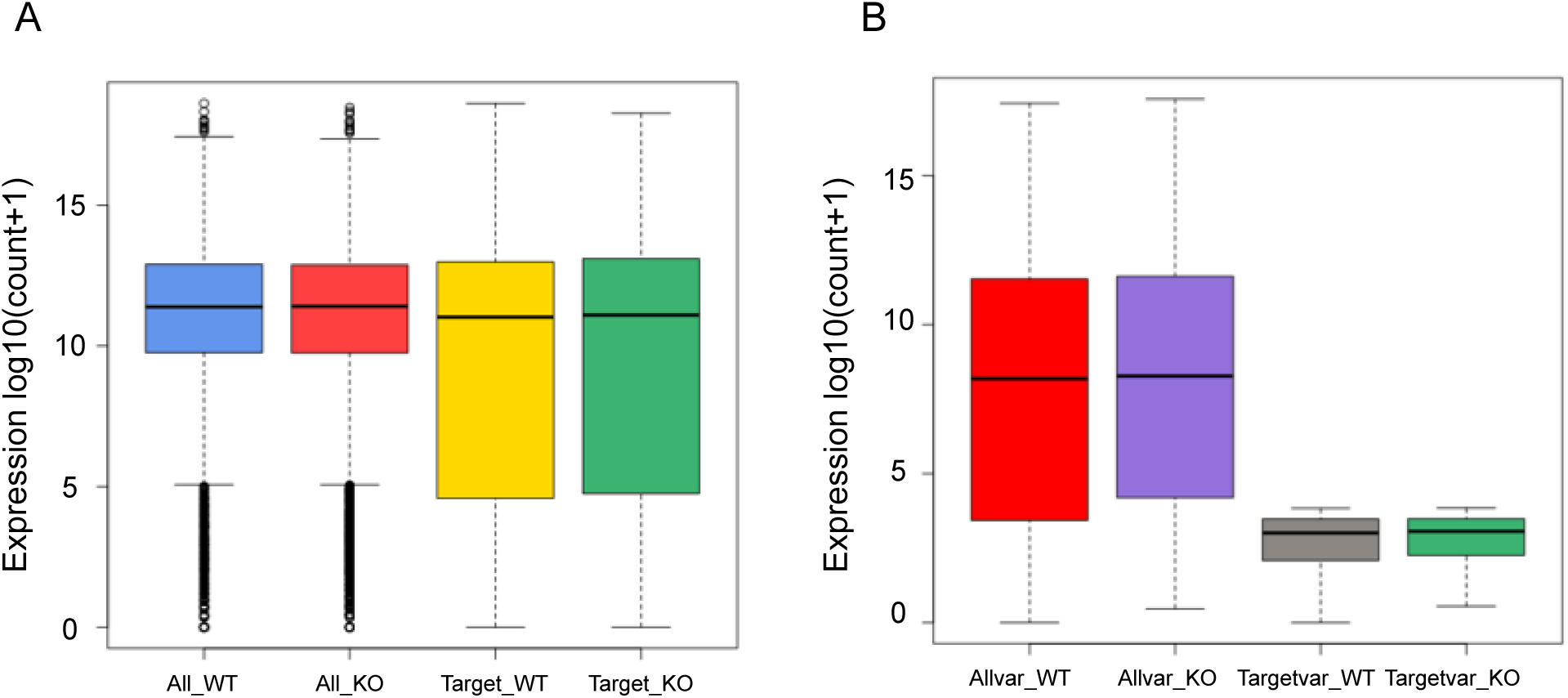
Effect of PfCDP1 knockout on total transcriptome and *var* genes. **(A)** The PfCDP knockout does not cause changes to the expression of a vast majority of parasite genes. The boxplots representing the average expression profile of all genes vs PfCDP target genes in PfCDP wildtype vs knockout condition. **(B)** The PfCDP knock out does not alter the expression of *var* genes, the boxplots representing the average expression profile of all *var* genes vs PfCDP1 target var genes in PfCDP1 wildtype vs knockout condition. The plots are for genes with >=1.8-fold PfCDP1 enrichment wherever relevant.

**Supplementary figure 11:**
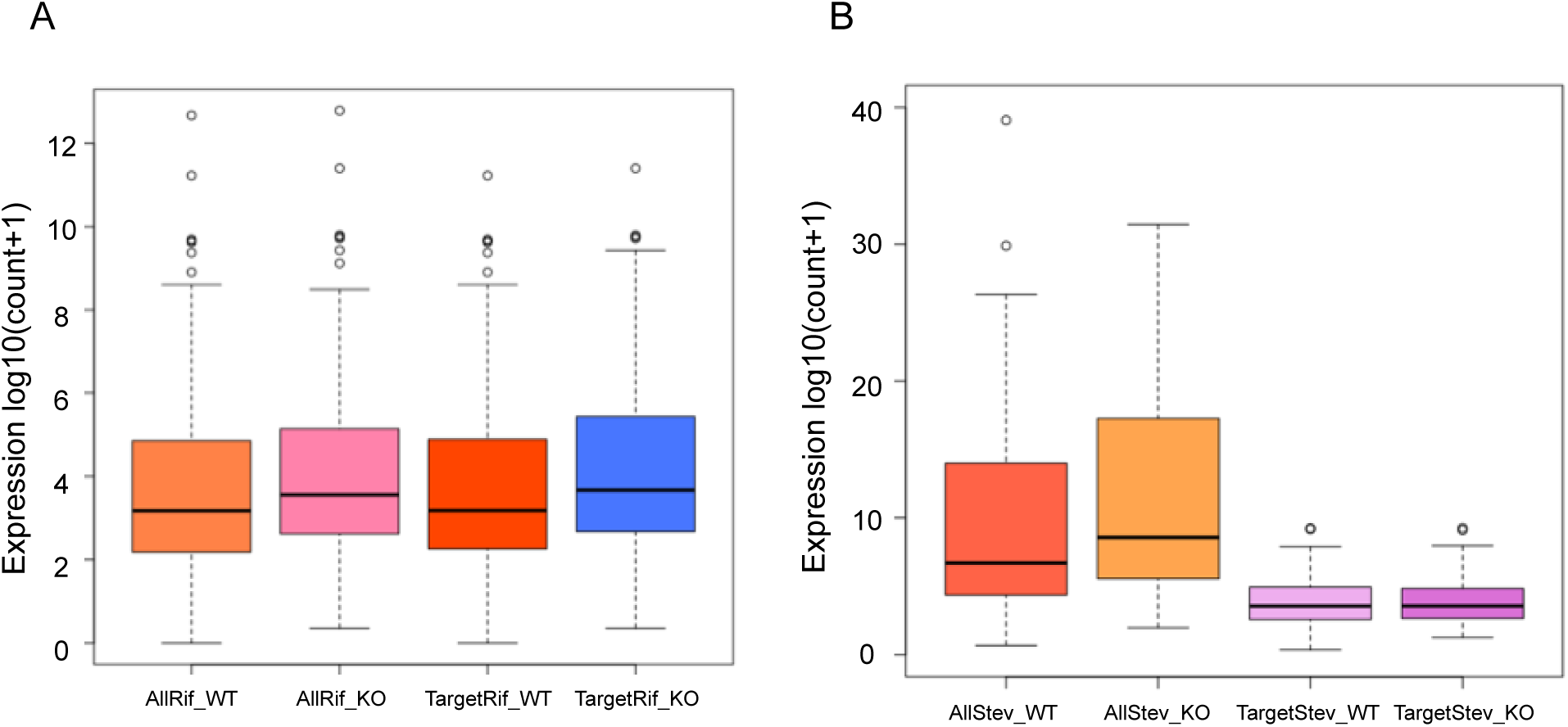
The PfCDP knockout causes changes in the expression of most *rifin* and *stevor* genes. **(A)** The boxplots representing the average expression profile of all *rifin* genes vs PfCDP target *rifin* genes in PfCDP wildtype vs knockout condition. Since a vast majority of rifin are targets of PfCDP the average expression of all rifins in these boxplots seen skewed with PfCDP1 knockout. **(B)** Boxplots representing the average expression profile of all stevor genes vs PfCDP1 target stevor genes in PfCDP1 wildtype vs knockout condition. The plots are for genes with >=1.8-fold PfCDP1 enrichment wherever relevant.

**Supplementary Figure 12:**
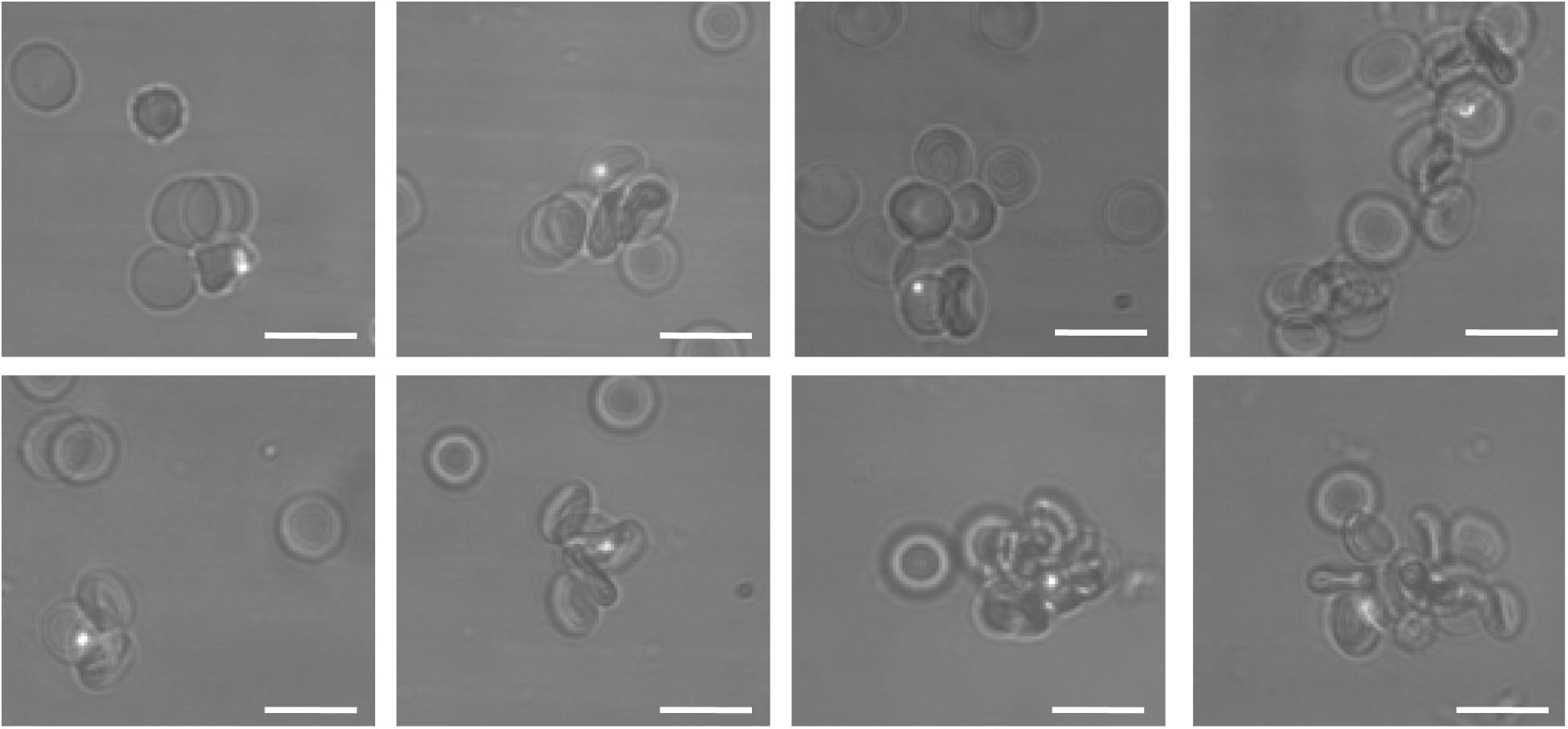
Images of rosette RBCs in PfΔCDP *P. falciparum* lines. The scale bar represents the 10 µm.

**Supplementary Table 1:**
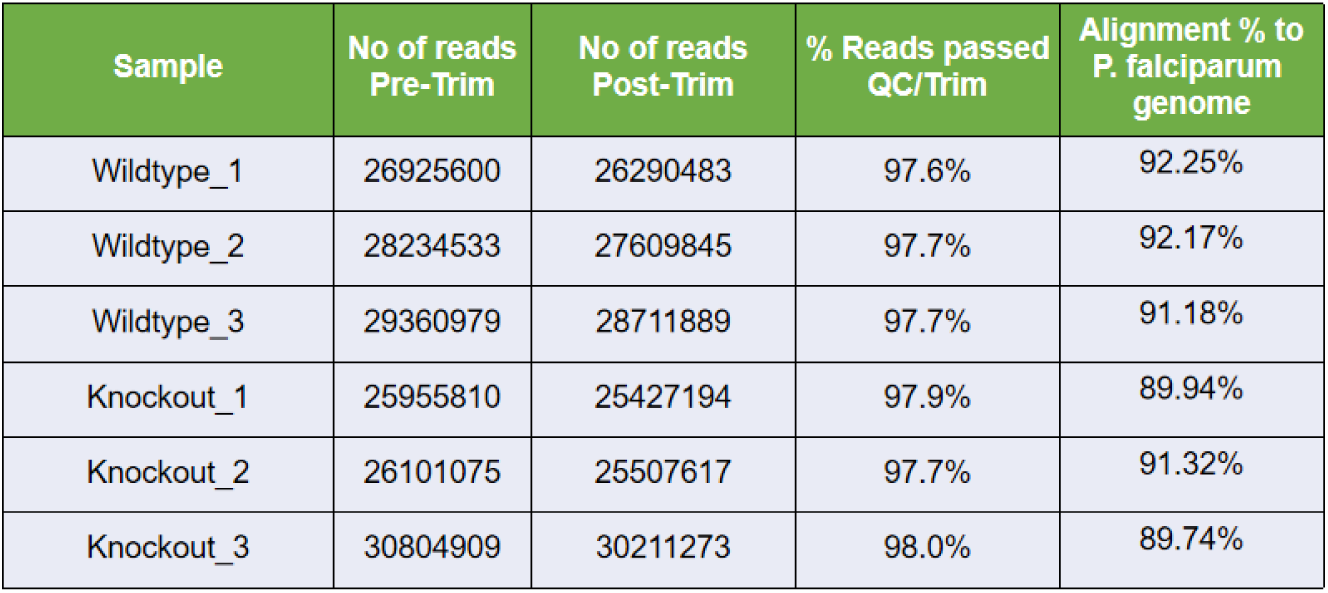
Quality Check/Trimming and target genome alignment statistics

| Sample | No of reads Pre-Trim | No of reads Post-Trim | % Reads passed QC/Trim | Alignment % to *P. falciparum* genome |
| --- | --- | --- | --- | --- |
| Wildtype_1 | 26925600 | 26290483 | 97.6% | 92.25% |
| Wildtype_2 | 28234533 | 27609845 | 97.7% | 92.17% |
| Wildtype_3 | 29360979 | 28711889 | 97.7% | 91.18% |
| Knockout_1 | 25955810 | 25427194 | 97.9% | 89.94% |
| Knockout_2 | 26101075 | 25507617 | 97.7% | 91.32% |
| Knockout_3 | 30804909 | 30211273 | 98.0% | 89.74% |

**Supplementary Table 2:**
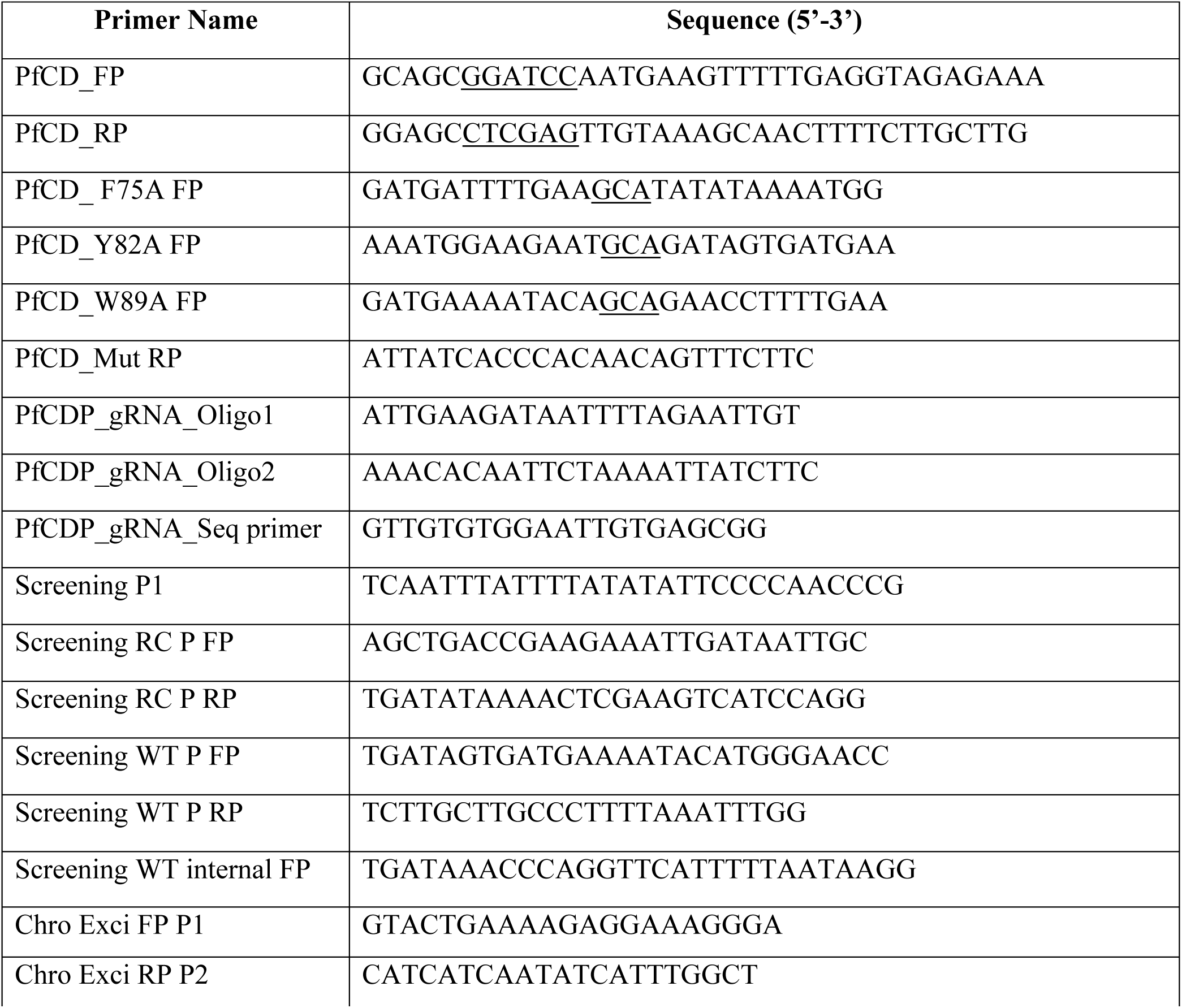
List of primers used in this study. The restriction sites for cloning into pGEX6P2 vector and mutated sites are underlined.

**Supplementary Table 3:**
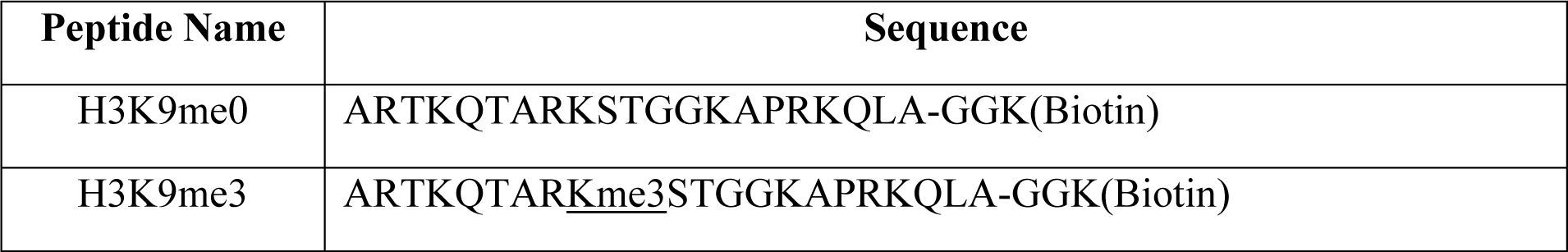
List of synthetic histone peptides used in this study. The target methylated amino acid in the peptide is underlined.

| Peptide Name | Sequence |
| --- | --- |
| H3K9me0 | ARTKQTARKSTGGKAPRKQLA-GGK(Biotin) |
| H3K9me3 | ARTKQTARK<u>me3</u>STGGKAPRKQLA-GGK(Biotin) |

